# Selecting genes for analysis using historically contingent progress: from RNA changes to protein-protein interactions

**DOI:** 10.1101/2024.05.01.592119

**Authors:** Farhaan Lalit, Antony M Jose

**Affiliations:** University of Maryland, College Park, MD, USA

**Author notes:** **Author Contributions:** A. M. J. designed the study; F. L. and A. M. J. performed the analyses (F. L. - all work on data tables and A. M. J. - all work on AlphaFold-based analyses); and F. L. and A. M. J. wrote the paper. **Competing Interest Statement:** The authors declare no competing interests.

**Keywords:** Mutual Information, AlphaFold, RNA silencing, homeostasis, *C. elegans*

## Abstract

Progress in biology has generated numerous lists of genes that share some property. But advancing from these lists of genes to understanding their roles is slow and unsystematic. Here we use RNA silencing in *C. elegans* to illustrate an approach for prioritizing genes for detailed study given limited resources. The partially subjective relationships between genes forged by both deduced functional relatedness and biased progress in the field was captured as mutual information and used to cluster genes that were frequently identified yet remain understudied. Some proteins encoded by these understudied genes are predicted to physically interact with known regulators of RNA silencing, suggesting feedback regulation. Predicted interactions with proteins that act in other processes and the clustering of studied genes among the most frequently perturbed suggest regulatory links connecting RNA silencing to other processes like the cell cycle and asymmetric cell division. Thus, among the gene products altered when a process is perturbed could be regulators of that process acting to restore homeostasis, which provides a way to use RNA sequencing to identify candidate protein-protein interactions. Together, the analysis of perturbed transcripts and potential interactions of the proteins they encode could help prioritize candidate regulators of any process.

## Introduction

Genes and gene products are often collected as lists based on unifying characteristics or based on experiments. Examples include genes that show enrichment of a chromatin modification, mRNAs that change abundance in response to a mutation, and proteins that interact with another protein. After the initial identification of a set of genes as belonging to a list, multiple approaches (1) are needed to generate an explanatory model. However, many genes do not receive further attention, as evidenced by recent meta-analyses, which highlighted numerous understudied genes in humans (2,3). Since single publications often analyze only one or a few genes, a wider view of genes with roles in a process could be gained by comparing lists generated by several studies. Such exploration could identify genes that are present in multiple lists but have not yet been selected for detailed study. Identifying these understudied genes is especially useful during the early stages of a field, when coherent models for most observed phenomena have not yet emerged. While this approach is also extensible to lists of anything that is used to characterize living systems (changes in lipids, metabolites, localizations, etc.), here we focus on lists of mRNAs, proteins, and small RNAs generated by the field of RNA silencing in the nematode *C. elegans*.

A gene present in many lists could be regulated in multiple separable ways and/or be regulated in one or a few ways by connected sets of regulators (Fig. 1A). For example, mRNA levels could be regulated through changes in transcription, turnover, localization, small RNA production, etc. or all changes could occur because of turnover regulation by a connected set of regulators. Changes in such genes could alter specific regulatory outputs, making them integrators of inputs from many other regulators. Alternatively, they could have no measurable consequence but might still be experimentally useful as general indicators of perturbation. One way that organisms could use general sensing of perturbation in a process could be to return the process to the pre-perturbation state through feedback (4). Such active resetting would enable restoration of homeostasis faster than through the dissipation of the perturbation alone.

**Figure 1.**
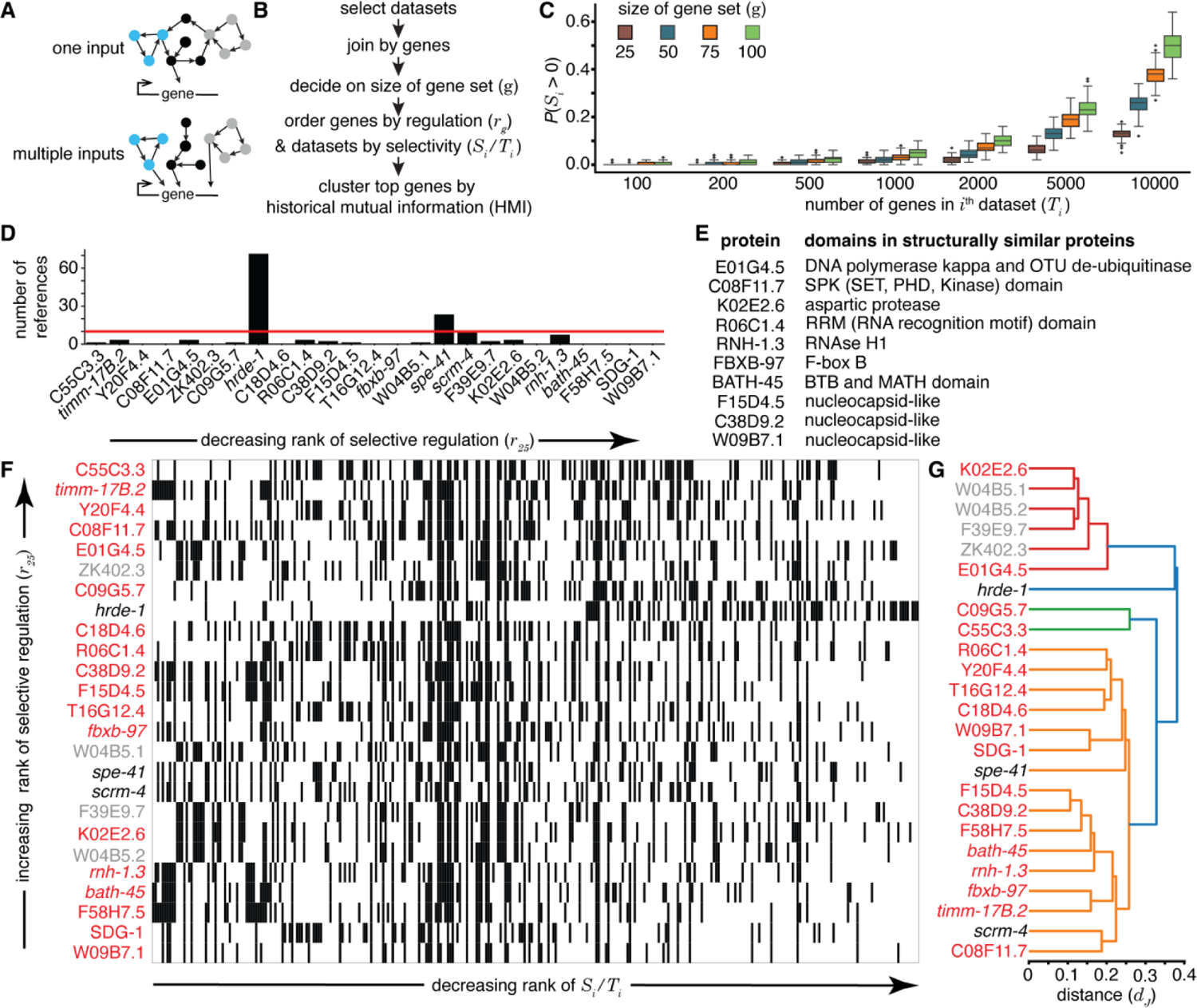
Some genes are selectively regulated, reported as part of many lists, and yet are understudied. (*A*) Schematics of possible regulatory architectures for genes found on multiple lists. (*top*) Gene receiving one input form a large network. (*bottom*) Gene receiving multiple inputs from separable networks. (*B*) Strategy for the identification of regulated genes. See Methods for details. (*C*) Relationship between *S_i_*, *T_i_*, and *g* obtained using simulated data for an organism with 20,000 genes. Distributions of the probabilities of having at least one overlapping gene within the selected gene set (*P*(*S_i_* > 0)) for 100 runs of each parameter combination are presented as box and whisker plots. (*D*) Numbers of publications listed on WormBase for the top 25 regulated genes ordered using *r_25_* in the field of RNA silencing in *C. elegans*. Red line marks 10 publications. (*E*) Domains present in proteins encoded by understudied genes among the top 25 genes that are suggestive of function. Proteins with high-confidence AlphaFold structures (12) were used to identify similar proteins as detected by Foldseek (17) or based on the literature ((18); C38D9.2, F15D4.5, and W09B7.2). (*F*) Heatmap showing the top 25 regulated genes. Presence (black) or absence (white) of each gene in each dataset is indicated. Relatively understudied (<10 references on WormBase) genes (red) or pseudogenes (grey) identified in (*D*) are indicated. (*G*) Hierarchical clustering of the top 25 genes based on co-occurrence in lists, where gene names colored as in (*F*) and ‘distance (*d_J_*)’ indicates Jaccard distance.

Here we present an approach to identify regulated but understudied genes in the field of RNA silencing in *C. elegans*. While these genes could play a variety of roles, we find that some of these genes encode predicted influencers of RNA-regulated expression that can directly interact with key regulators of RNA silencing. Others could serve as regulatory links that connect RNA silencing with other processes. The many hypotheses generated through these analyses need to be evaluated using experiments that selectively perturb regulatory interactions.

## Materials and Methods

Data tables from 82 studies on RNA silencing in *C. elegans* that were published between 2007 and 2022 were downloaded (Table S1), reformatted manually and/or using custom scripts, and filtered to generate lists that only include entries with reported p-values or adjusted p-values < 0.05, when such values were available. Gene names were standardized across datasets using tools from Wormbase (5). The top ‘*g*’ genes that occur in the greatest numbers of tables were culled as the most frequently identified genes. A measure for the extent of regulation of each gene (*r_g_*) was used to aid their prioritization for detailed study. Co-occurrence patterns of genes in different tables were captured using the Jaccard distance (*d_J_*) (6) or a symmetric measure of normalized mutual information (7), defined here as Historical Mutual Information (HMI). The *d_J_* values were used to generate a dendrogram using the average linkage method (Fig. 1). HMI was used to group genes into clusters according to the Girvan-Newman algorithm (8) and different sets of genes were highlighted (Fig. 8). Gene ontology (GO) analyses were performed using Gene Ontology Resource (https://geneontology.org/; (9,10)).

Prediction of dimer formation between the 18 proteins encoded by understudied genes among the top 25 genes and 25 key regulators of RNA silencing were obtained using AlphaFold 2 (11,12) run on a high-performance cluster (Zaratan at UMD) and/or using the AlphaFold 3 (13) server online (https://golgi.sandbox.google.com/). Large regulators (DCR-1, EGO-1, ZNFX-1, NRDE-2, and MET-2) were tested on the AlphaFold 3 server initially and positive hits, if any, were examined again using AlphaFold 2 (e.g., interaction of EGO-1 with W09B7.1). The computed models were processed using custom shell scripts, python programs, and ChimeraX (14). Briefly, the highest ranked model for each pair of proteins was depicted with the predicted aligned error used to highlight inter-protein interactions as pseudobonds colored according to the alphafold pae palette on ChimeraX (Movie S1 to S93). For models that satisfy the criteria for maxPae (<5 Å) and for distance (<6 Å), an approximation of the interaction area was calculated by isolating the mutually constrained residues and using the ‘buriedarea’ command (ChimeraX). This area was divided by the product of the number of amino acids in each protein to get a normalized value and scaled uniformly before plotting (e.g., Fig. 2B). Finally, the ranking scores (0.8*ipTM + 0.2*pTM for AlphaFold 2.3 and 0.8*ipTM + 0.2*pTM + 0.5*disorder for AlphaFold 3) were used to shade the circle representing each interaction (Fig. 2B) and/or plotted (Fig. 5A). All interactions predicted in the study were summarized into a network diagram (Fig. 8G) using Gephi and Adobe Illustrator. See Supplemental Material for detailed materials and methods.

**Figure 2.**
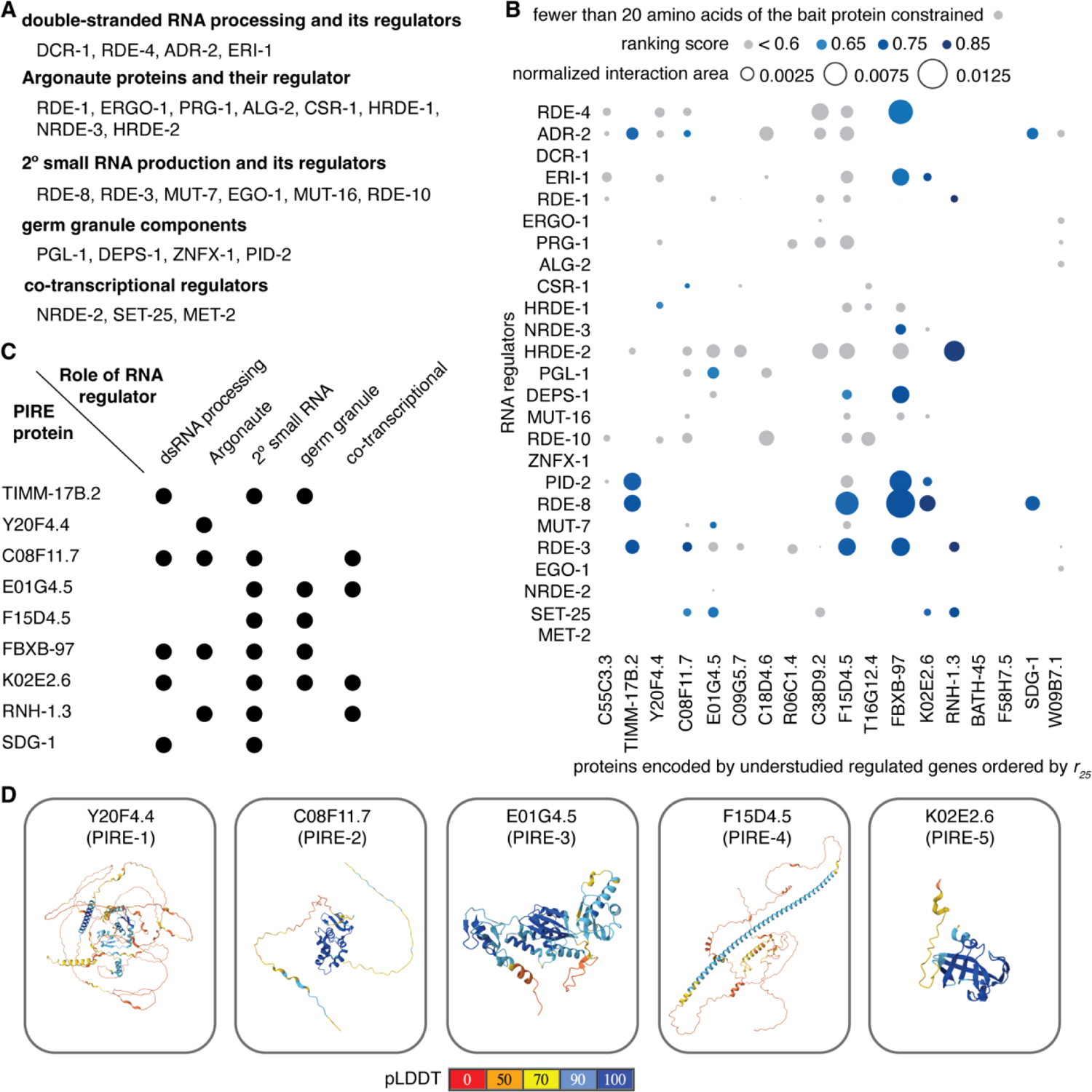
Understudied regulated genes encode proteins predicted to interact with key regulators of RNA silencing. (*A*) Regulators of RNA silencing in different categories examined for predicted interactions with proteins encoded by understudied genes identified in this study. See text for details. (*B*) Predicted interactions between proteins encoded by top 25 genes ordered by their *r_25_* scores and known regulators of RNA silencing in *C. elegans*. The area of the interaction surface between partners normalized by the product of the sizes of the interactors is shown as a bubble plot (inter-protein predicted aligned error <5Å and inter-residue distance <6Å). Interactions with a low ranking score (< 0.6) and/or that constrain fewer that 20 amino acids in proteins encoded by the understudied genes are indicated in grey. Also see Fig. S1 and Movies S1 to S32. (*C*) Proteins encoded by understudied genes with significant interactions are predicted to impact multiple steps in RNA silencing. (*D*) Predicted structures for the five newly named predicted influencers of RNA-regulated expression (PIRE) proteins are shown with the per-residue confidence (pLDDT) as present in the AlphaFold protein database (83).

## Results

### Many genes have been repeatedly reported within data tables but remain understudied

To determine if there are any understudied regulated genes that are relevant for RNA silencing in *C. elegans*, we examined data from past studies in the field. While complete replication of each study might be needed for direct comparisons, this goal is impractical. Even beginning with the ‘raw’ data deposited to public resources (e.g., fastq files after RNA-seq) and repeating the analyses reported in a publication is not always feasible. Summary tables from previous analyses presented in publications provide a practical intermediate level of data to use for comparisons across studies. Therefore, we collated a total of 398 tables from 82 publications for comparison (see methods and Table S1 for list of tables) and joined the tables together after standardizing gene names to yield genes that can be compared for presence or absence across the 398 lists (Fig. 1B). About 86% (342 of 398) of the included gene lists document RNA changes (mRNA, small RNA, or total RNA) that accompany a perturbation. Of the remaining ∼14%, some lists document co-immunoprecipitating proteins (19), were pre-defined enrichment lists (24), or are based on other experiments (13). To prioritize a set of genes (*g*) that receive extensive regulatory input and/or that encode proteins that interact with many other proteins and are yet included in selective lists, we propose a metric *r_g_* (Fig. 1B). Since the likelihood of including a gene from the lists increases with *g*, the metric is specified with a subscript for each analysis (e.g., *r*_25_ refers to a regulation score when the top 25 genes that are most commonly present in lists are considered) and defined to be:

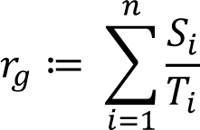

where *g* = size of gene set chosen for analysis, *n* = total number of lists with altered genes, *S*_*i*_ = number of genes from the *i*^th^ list that is also present in the gene set *g*, and *T*_*i*_ = total number of genes in the *i*^th^ list. The larger the set of genes (*g*) selected, the greater the chance of a dataset (with *T_i_* genes) having at least one overlapping gene within the selected gene set (probability given by *P*(*S_i_* > 0) in Fig. 1C). The metric *r*_*g*_ is a decision aid that helps with choosing genes for experimental analysis and is not to be taken as an objective measure of the importance of the gene for the biological process under study.

The top 25 genes sorted according to their *r*_25_ values included the germline Argonaute HRDE-1 (15), which has been the subject of numerous studies (Fig. 1D). While most other genes are understudied (fewer than 10 publications on WormBase), among the 25 genes is *sdg-1*, which was recently reported to be regulated by the double-stranded RNA (dsRNA) importer SID-1 and encodes a protein with a suggested role in feedback regulation of heritable RNA silencing by colocalizing with perinuclear germ granules (16). This discovery suggests that the analysis of the additional genes with high *r*_25_ values could also be fruitful. Of the 18 understudied genes that encode proteins, seven had predicted structures of high confidence (i.e., domains with predicted local distance difference test (pLDDT) > 90) in the AlphaFold Protein Structure Database (12). These structures were then used to identify related protein domains using Foldseek (17) (Fig. 1E; E-value < 0.05). These include conserved domains, most of which have known biochemical activities: de-ubiquitinase (E01G4.5), SPK (C08F11.7), aspartic protease (K02E2.6), RNAse H1 (RNH-1.3), F-box B (FBXB-97), BTB plus MATH (BATH-45), and RNA Recognition Motif (R06C1.4). Three more proteins have been proposed to be nucleocapsid-like proteins encoded by genes within retrotransposons ((16,18); F15D4.5, C38D9.2, and W09B7.1 in Fig. 1E). These candidates can be experimentally analyzed in the future for possible roles in RNA silencing. To explore the relationships between these genes (Fig. 1*F* and 1*G*), we clustered the genes and generated a dendrogram where genes present together in different lists are closer together (see supplementary methods). The dendrogram had a cluster (red in Fig. 1G) that included all four pseudogenes, suggesting that this method could capture functional relatedness despite the limitations and biases introduced by the available data.

### Multiple proteins encoded by the top 25 genes are predicted to interact with known regulators of RNA silencing

In general, understudied regulated genes could play diverse roles, some of which could impact RNA silencing. Such feedback during RNA silencing is supported by recent observations. For example, animals typically recover from silencing initiated by dsRNA within the germline (19) or in somatic cells (20). This recovery occurs despite the presence of amplification mechanisms, suggesting that silencing ends either when the trigger dsRNA runs out and/or because of homeostatic control through feedback inhibition. In support of self-limiting behavior that is expected upon feedback inhibition, an inhibitor of RNA silencing is recruited to genes targeted by dsRNA (21) and a regulatory loop limits the production of some endogenous small RNAs (22). Some of the top understudied regulated genes identified here could play a role in the homeostatic return after perturbation. The return could be achieved by modulating the activity of factors that promote RNA silencing in a variety of ways, including regulation of transcription, post-transcriptional RNA processing, RNA localization, translation, post-translational modifications on proteins, protein localization, etc. Of these possibilities, one that could be surveyed computationally is regulation through direct protein-protein interactions. Therefore, to test if any of the proteins encoded by the top 25 genes could interact with known regulators of RNA silencing, we examined the potential for protein-protein interactions using their predicted structures.

We selected 25 known regulators of RNA silencing (see Fig. 2A) chosen for their roles in different phases of the deduced mechanism(s) of RNA silencing (23–25). These include proteins with roles in the processing of dsRNA and its regulators; Argonaute proteins and their regulators; proteins with roles in secondary small RNA production and its regulators; components of germ granules; and co-transcriptional regulators (Fig. 2A). We then examined their predicted interactions with the 18 proteins encoded by understudied regulated genes among the top 25 (highlighted in red, Fig. 1G). For 20 RNA regulators, we used AlphaFold 2, which makes extensive use of multiple sequence alignments for computing inter-protein interactions and has a success rate of ∼50-60% (26,27). Since the computational cost of AlphaFold 2 escalates with the number of amino acids, interactions with the remaining 5 larger regulators (DCR-1, EGO-1, ZNFX-1, NRDE-2, and MET-2) were tested on the recently available but proprietary AlphaFold 3 server (13), which can predict interactions with ligands, and as with AlphaFold 2, uses multiple sequence alignments for its structure predictions. To stratify the predicted interactions, we initially considered the maximal inter-protein predicted aligned error (PAE) and the distance between the interacting residues (distance), which was allowed to be up to twice the length of hydrogen bonds (∼3 Å (28)). Examining interactions with a criterion of PAE less than 5 (which is more stringent than the 8 Å error that has been used successfully (29)) revealed numerous interactions (blue in Fig. 2B). Therefore, to constrain the predictions further, we used the ranking scores, which are a combination of interface-predicted template modeling (ipTM) and predicted template modeling (pTM) scores: 0.8*ipTM + 0.2*pTM for AlphaFold 2.3 (11) and 0.8*ipTM + 0.2*pTM + 0.5*disorder for AlphaFold 3 (13). We only considered interactions with a ranking score greater than 0.6, which is relatively high given than ipTM scores as low as ∼0.3 can yield true positives (30), and that constrain a minimum of 20 residues in the proteins encoded by understudied genes (not grey in Fig. 2B), which we define as predicted interactions of high confidence. Together, these criteria identified 32 interactions (Fig. 2B and Movies S1 to S32). Among the regulators, RDE-3 and RDE-8 had the highest numbers of predicted interactors (5 proteins each) and among the proteins encoded by understudied genes, FBXB-97 had the highest number of predicted interactors (7 proteins). These high-confidence interactors included proteins that were predicted to interact with every phase of the deduced mechanism(s) for RNA silencing (Fig. 2C). Since the precise numbers of interacting residues required for a meaningful interaction in vivo is variable and unknown, interactions that constrain fewer residues could have measurable impacts on function. Nevertheless, we conservatively designate each protein that is predicted to interact with one or more RNA regulators with relatively high confidence as a **P**redicted **I**nfluencer of **R**NA-regulated **E**xpression (PIRE). We refer to five of these as PIRE-1 through PIRE-5 (Y20F4.4, C08F11.7, E01G4.5, F15D4.5, and K02E2.6, respectively; Fig. 2D) and preserve the names of the four that were already given names based on structural homology (subunit of the **T**ranslocase of the **I**nner **M**itochondrial **M**embrane TIMM-17B.2, the **F**-**b**o**x B** protein FBXB-97 and the **RN**ase **H** protein RNH-1.3) or after detailed study (the **S**ID-1-**d**ependent **g**ene protein SDG-1). For convenience, these nine putative interactors (and any additional interactors identified below) are collectively referred as PIRE proteins here. This provisional designation can be amended should more specific information regarding their roles be obtained through future experimental studies.

Each of these interactions (Fig. 2B and Movie S1 to S32) suggest hypotheses for their functional impact based on the known roles of RNA regulators and the domains present in PIRE proteins (Fig. 1E). The two PIRE proteins encoded by genes within retrotransposons (PIRE-4 and SDG-1) that also interact with regulators of RNA silencing, supports the idea that retrotransposon-encoded genes influence their own RNA-mediated regulation (e.g., (16)). FBXB-97, which is predicted to be an F-box protein (31), could promote ubiquitin-mediated degradation of its interactors (RDE-4, ERI-1, NRDE-3, DEPS-1, PID-2, RDE-8, and RDE-3), sequester them (inhibiting their activity), or potentially promote ubiquitination of their interacting RNA (32). A precedent for such an intersection between ubiquitin-mediated protein degradation and small RNA-mediated RNA regulation is the role for a ubiquitin ligase in degrading Argonautes when there is extensive base-pairing between miRNAs and their targets (33,34). PIRE-5, which is predicted to be a protease, could cleave its interactors (ERI-1, PID-2, RDE-8, and SET-25) to regulate their activity – a mode of regulation that has been recently elucidated for Argonaute proteins (35) and implicated in RNA silencing within the germline (36). Additional PIRE proteins with confidently predicted domains (e.g., RNase H in RNH-1.3, multiple domains in PIRE-3, and SPK domain in PIRE-2) potentially implicate new biochemical activities in the process of RNA silencing. In all, two general modes of interaction between PIRE proteins and the tested regulators of RNA silencing that are not mutually exclusive could be discerned (Fig. 3). In one mode exemplified by FBXB-97 (Fig. 3, *left*), the interactions with most regulators involve nearly the same set of residues. In the other mode exemplified by PIRE-3 (Fig. 3, *right*), interactions with different regulators involve different sets of residues. In summary, predictions using AlphaFold identify numerous interactions that inspire follow-up work to test hypotheses about the roles of PIRE proteins in RNA silencing.

**Figure 3.**
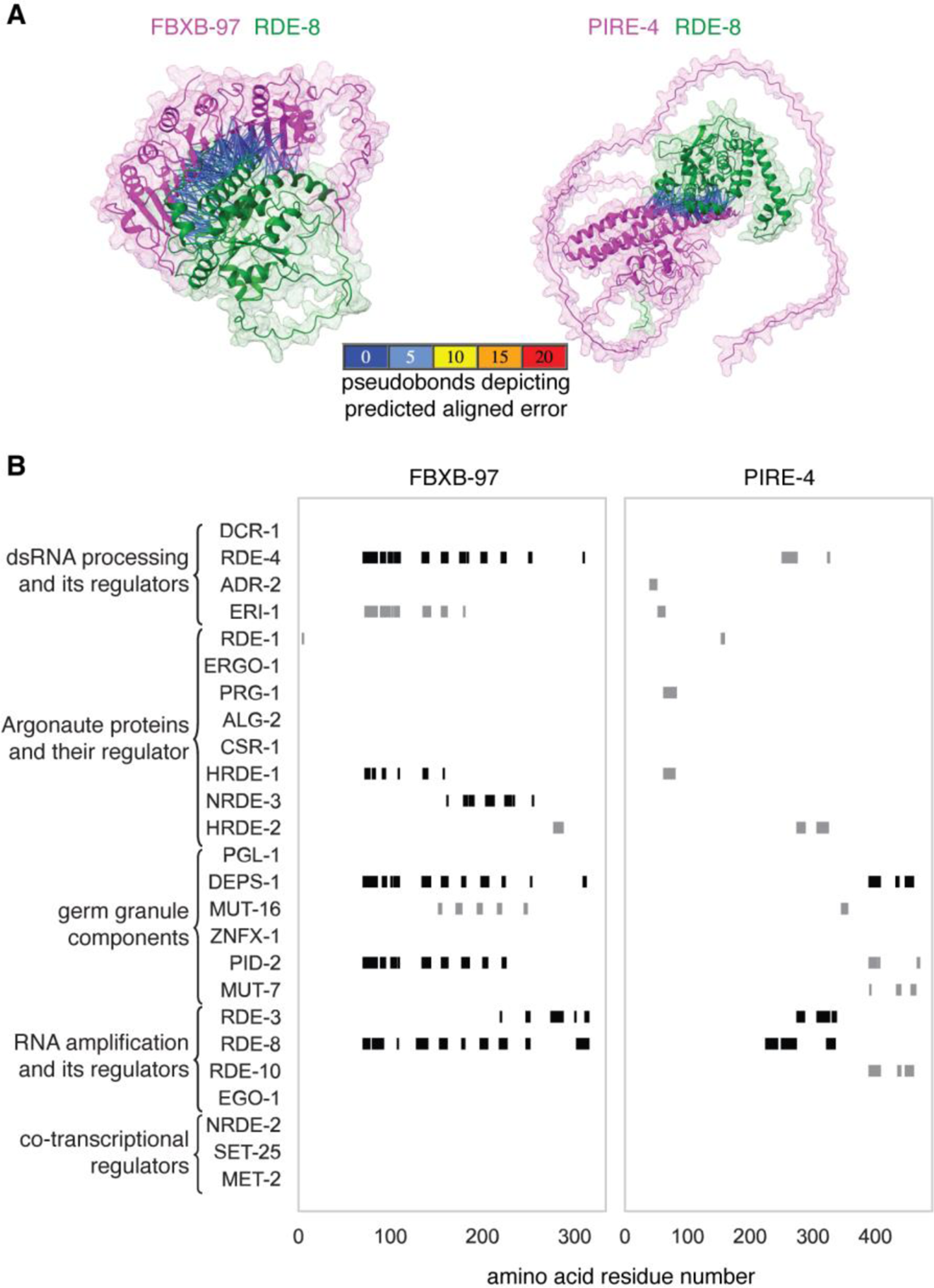
Predicted Influencer of RNA-regulated Expression (PIRE) proteins interact with regulators of RNA silencing in two general modes. (*A*) Predicted interactions between the PIRE proteins (magenta) FBXB-97 (left) and PIRE-3 (right) with the known regulator RDE-8 (green) that are of high confidence (constraining more than 20 amino acid residues with an inter-Cα distance less than 6Å and PAE less than 5Å) are indicated with pseudo bonds. (*B*) Regions of the PIRE protein sequence constrained by the interacting regulator. Markers (black, ranking score >0.6; grey, ranking score <0.6) are enlarged with respect to the X-axis for visibility (e.g., the marker denoting the interaction between RDE-1 and FBXB-97 only indicates one residue).

### Predictions by AlphaFold 2 and AlphaFold 3 do not always agree

While AlphaFold 2 predicted all the interactions classified as high-confidence interactions, the one interaction predicted by AlphaFold 3 (EGO-1 and W09B7.1) with a maximal PAE <5Å and distance <6Å constrained fewer than 20 residues (Fig. 2B, Fig. S1, and Fig. S2). The reason for this extreme discrepancy is unclear.

To directly compare both approaches for predicting protein-protein interactions, we examined some of the interactions predicted by each approach using the other. We first examined significant interactions predicted with a high ranking score according to AlphaFold 2 (> 0.8, Fig. 4A). Of these, only the interaction between RNH-1.3 and RDE-3 was confidently predicted by AlphaFold 3, albeit with a lower score (0.85 for AF2 vs 0.68 for AF3). Aligning both predicted complexes using the RDE-3 protein revealed that both predictions are in good agreement (Fig. 4B and Movie S33). We next considered two proteins, FBXB-97 and PIRE-4, for which multiple interactors were predicted by AlphaFold 2 with varying confidence. While AlphaFold 2 predicted interactions between PIRE-4 and 3 RNA regulators (ranking = 0.73, 0.70, and 0.61), and between FBXB-97 and 7 RNA regulators (ranking = 0.66, 0.67, 0.75, 0.78, 0.75, 0.77, and 0.76), AlphaFold 3 only predicted an interaction with RDE-3 for both proteins (Fig. 4C, ranking = 0.78 and 0.79). The RDE-3-interacting residues of FBXB-97 predicted by both approaches overlapped but those of PIRE-4 did not (Fig. 4C). Furthermore, aligning the predicted protein-protein complexes using RDE-3 showed a large discrepancy in the positions of the interacting partners in both cases (Fig. 4D, FBXB-97, *left*; PIRE-4, *right*; and Movies S34 and S35). Similarly, comparing the predictions for interactions between EGO-1 and W09B7.1 also revealed large discrepancies (Fig. 4E and Movie S36). While a region of interaction was predicted using AlphaFold 3 with PAE <5Å and distance <6Å (Fig. 4E, right), regions of interaction were only detectable using AlphaFold 2 when the maximal PAE allowed was increased to 10 (Fig. 4E, left). Even at this lower threshold for error, the predicted interacting regions differed between the two approaches (black ovals in Fig. 4E).

**Figure 4.**
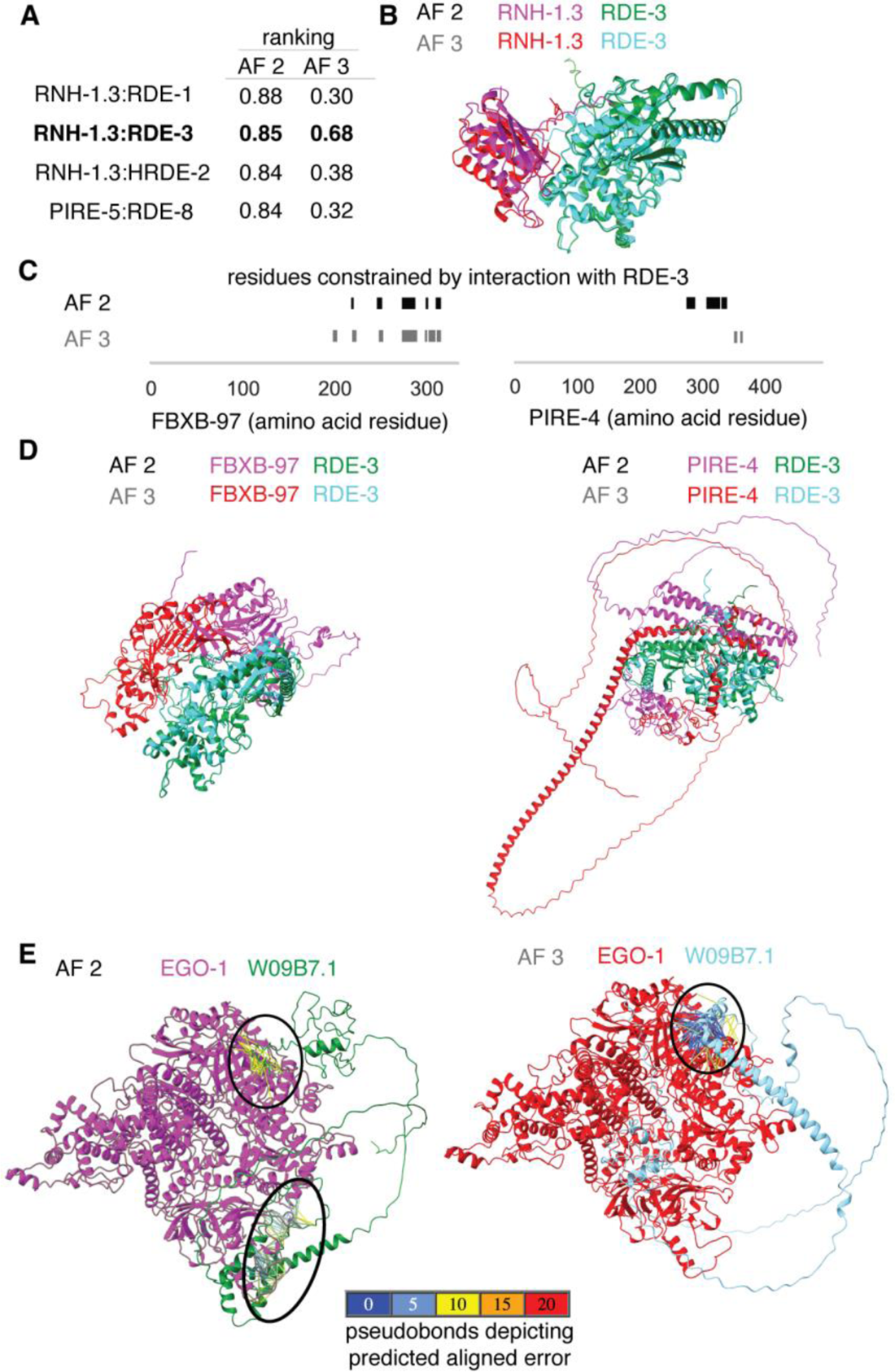
Interactions predicted by AlphaFold 2 and by the AlphaFold 3 server can differ. (*A*) Comparison of the top ranking interactions between known regulators of RNA silencing and the PIRE proteins predicted by AlphaFold 2 (AF 2 (11); 0.8*ipTM + 0.2*pTM) with the score generated by AlphaFold 3 (AF3 (13); 0.8*ipTM + 0.2*pTM + 0.5*disorder). A high-confidence prediction by both approaches is highlighted in bold. (*B*) Models for the interaction of RNH-1.3 with RDE-3 generated by AF2 and AF3 overlayed using RDE-3. Also see Movie S33. (*C*) Comparison of residues of PIRE proteins constrained through interactions as predicted by AF2 (black) or by AF3 (grey). (*D*) Comparison of interactions between FBXB-97 and RDE-3 (left), and between PIRE-4 and RDE-3 (right) as predicted by AF2 (black) and the AF3 server (grey), respectively. Structures are shown with differential coloring of each protein and overlayed using the RDE-3 structures in both cases. Also see Movies S34 and S35. (*E*) Interactions between EGO-1 (magenta or red) and W09B7.1 (green or cyan) predicted by AF2 or AF3. Black ovals indicate interacting regions with inter-protein PAE <10Å (left) or <5Å (right). Also see Movie S36.

The reasons for the differences between predictions by AlphaFold 2 and AlphaFold 3 could be varied. For example, differences in sampling of predictions, which is expected to correlate with success rate (37) (25 models per AlphaFold 2 run versus five per AlphaFold 3 run on the server) and/or differences in handling intrinsically disordered regions, for which structures can be identified by AlphaFold 2 if they conditionally fold (38). Modifications to these algorithms that extend capabilities continue to be developed (e.g., modeling of interacting interfaces within intrinsically disordered regions (39), predicting multiple conformations (40), and predicting large protein assemblies (41)). Therefore, further comparisons as newer algorithms for predicting protein-protein interactions continue to be developed (e.g. (42)) and customized exploration of criteria for interactions (43) may be useful for determining when each algorithm can aid the generation of hypotheses.

### Convergence and rarity of models support the use of flexible criteria for initial screens

The RNH-1.3:RDE-3 complex is supported by relatively high-confidence models predicted using AF 2 and AF 3 (Fig. 4*A* and 4*B*), *rnh-1.3* RNA accumulates in *rde-3*(-) animals (44,45), and both *rnh-1.3* and *rde-3* were featured in the abstract of an early publication (46). Therefore, we examined the predicted interactions between RNH-1.3 and RDE-3 in detail to refine the criteria for identifying candidate interactors and to analyze the potential reasons for differences between predictions by AF 2 and AF 3.

While the high-confidence model of the RNH-1.3:RDE-3 complex by AF 2 had a ranking score of 0.85, all the other 24 models from the run had much lower scores (Fig. 5A). In fact, the highest-ranking score in 17 subsequent runs was only 0.51. This rarity of high-scoring models suggests that multiple runs may be required on the AlphaFold 3 server as well to discover the high scoring models. Consistently, only one of 5 new runs of AF 3 resulted in a high-scoring model (Fig. 5B). Interestingly, an overlay of models with the top two ranks showed a highly similar structure both in the case of AF 2 and AF 3 predictions despite the large differences in their scores (0.85 vs. 0.51 for AF 2 and 0.74 vs. 0.35 for AF 3 in Fig. 5C; and Movies S37 and S38). This observation suggests that for some protein complexes, the models predicted with relatively low ranking scores could nevertheless be close to the highest-scoring model. To systematically analyze the convergence of the models with ranking scores, we overlayed the highest scoring models from the 18 runs of AF 2 and calculated the root mean square deviation (RMSD) of each model from the highest-scoring one (Fig. 5D). Scores as low as ∼0.4 resulted in models that were within ∼4Å RMSD of the highest-scoring model. However, scores below that (red line in Fig. 5D) were associated with models that could either have a low (e.g., < 10Å) or high (e.g., > 20Å) RMSD compared with the highest-scoring model. These analyses suggest that models with a ranking score of 0.4 could be worth exploring further, although we have preserved the more conservative threshold of 0.6 in all subsequent analyses using AlphaFold 2. To make the contributions of the two interacting proteins symmetric, we propose that the product of the number of constrained residues in the bait (n_bait_) and that in the prey (n_prey_) be greater than 100. Using these revised criteria, we re-examined all AF 2 predictions (Fig. S3) and found that all 32 previously predicted interactions (Fig. 2B) were preserved, and an additional 10 interactions were predicted as significant (Fig. S3). These additional interactions (Movies S39 to S48) include those of two RNA regulators with R06C1.4 (designated PIRE-6), two RNA regulators with C38D9.2 (designated PIRE-7), and one RNA regulator with T16G12.4 (designated PIRE-8).

**Figure 5.**
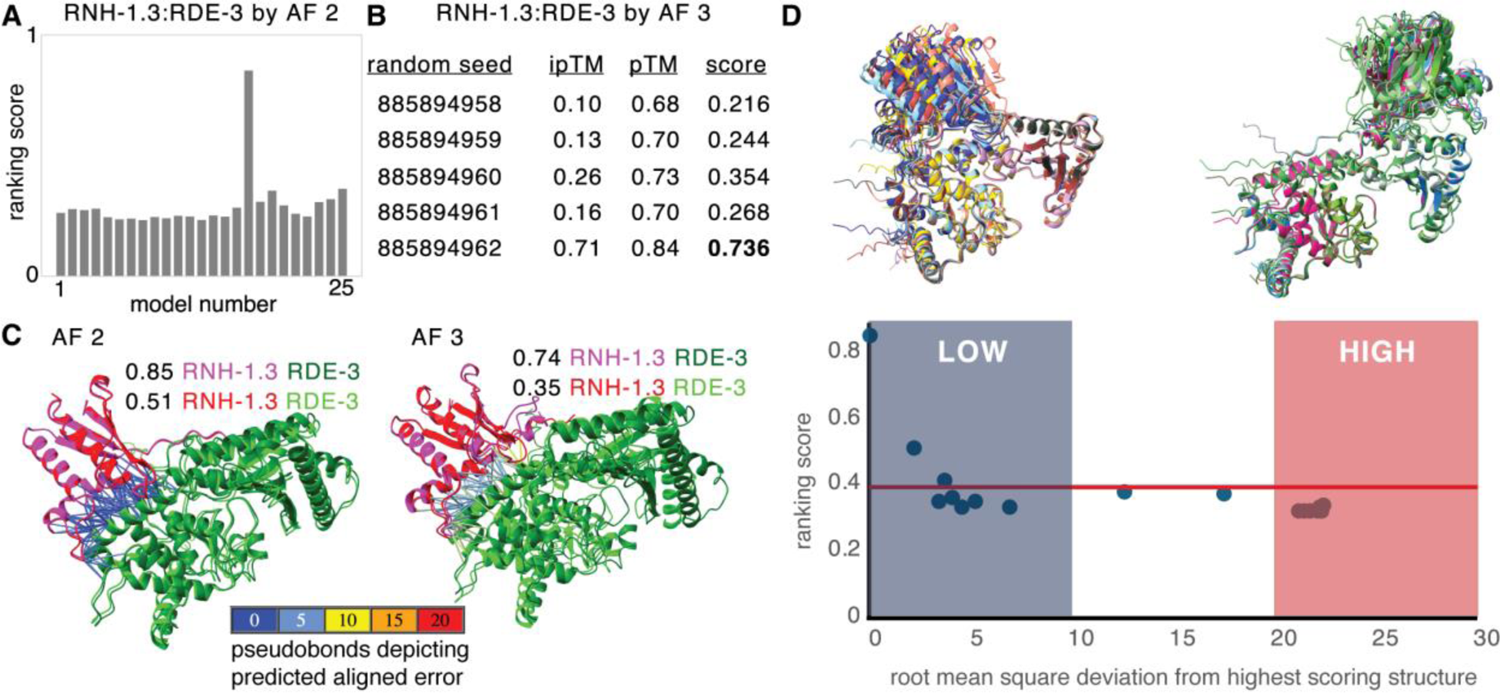
High-ranking models can be rare, and models can converge early with increasing scores. (*A*) Distribution of ranking scores for the 25 models of RNH-1.3:RDE-3 generated by AF 2. (*B*) Multiple runs with different random seeds and resulting scores for models of RNH-1.3:RDE-3 generated by AF 3. (*C*) Overlay of models with the highest scores from two different runs showing similar interactions between RNH-1.3 (magenta or red) and RDE-3 (green or lime) predicted by both AF 2 and AF 3. Pseudobonds depicting the predicted aligned errors for the constrained residues are highlighted for both pairs of models. Also see Movie S37 and S38. (*D*) A range of scores can underlie nearly similar architectures of a predicted complex. The highest scoring model for RNH-1.3:RDE-3 from each of 18 AF 2 runs (different colors) were superimposed using RDE-3. *Top*, Superimposed models for low (less than 10Å) and high (more than 20Å) root mean square deviation (RMSD) values are shown. *Bottom*, Ranking scores are plotted after arranging models in increasing order of RMSD from the highest scoring model.

Taken together, these analyses suggest heuristics for managing false negatives. Since high-scoring models can be rare (1 in 450 AF 2 models with a score > 0.6 for RNH-1.3:RDE-3) and therefore require many runs to discover, false negative rate can be set based on available computational resources. Given the early convergence of some models with increasing ranking score (a model with ∼0.4 ranking score only had an RMSD of ∼4Å compared with the highest scoring model for RNH-1.3:RDE-3), reducing the ranking threshold could lead to the discovery of interactors within fewer runs. In contrast, false positives are difficult to estimate or manage because we would need a set of proteins that would not interact with each other under any circumstance – such an idealized set may not exist.

### Pseudogenes among the top 25 genes could encode proteins that interact with some RNA regulators

Since pseudogenes could have the potential to encode peptides, we checked for this possibility in the four identified among the top 25 genes. Examination of all possible reading frames revealed uninterrupted stretches that could code for peptides for each ‘pseudogene’ (F09E9.7 - 141 aa, W04B5.1 - 85 aa, W04B5.2 - 144 aa, and ZK402.3 - 158 aa). These peptides if expressed have the potential to interact with some of the known RNA regulators tested (6 of 80 possible interactions in Fig. S4; Movies S49 to S54). These could reflect interactions between peptides from these ‘pseudogenes’ or from the corresponding coding genes. In support of this idea, the STAU-1-like peptide that could be encoded by the ‘pseudogene’ F39E9.7 and the dsRNA-binding protein STAU-1 (47) are both predicted to interact with ADR-2, CSR-1, and RDE-8 (Fig. S4; Movies S55 to S60). These results highlight the possibility that genes annotated as non-coding RNAs, or pseudogenes could have a role encoding a regulatory peptide.

### Multiple predicted interactors of RDE-3 suggest regulated production of poly-UG RNAs

Currently, information on the interactors of any protein in *C. elegans* is curated at WormBase (5) and the Alliance of Genome Resources (48) websites. We selected the poly-UG polymerase RDE-3, which catalyzes the production of a key intermediate of RNA silencing called poly-UG RNAs (45,49,50), as a case study to examine the value added by analysis using AlphaFold, if any. The websites list 5 physical interactors of RDE-3 identified through experiments reported in multiple publications (MUT-7(51), MUT-16(52), PIK-1(53), PRG-1(54), and RDE-8(55)). Including these putative direct interactors, we tested the interaction of 22 regulators of RNA silencing and found significant interactions with 12 proteins using AF 2 (Fig. 6A; Movies S61 to S72). Subsets of proteins appear to constrain different sets of residues on RDE-3 (Fig. 6B), suggesting different consequences on RDE-3 activity upon interaction for different groups of proteins. Taken together with the previously discovered interactions, a total of 19 interactors are predicted for RDE-3 by AF 2 (Fig. 6C). Of these, only 6 interactors were also identified by AF 3 when searched using 5 different random seeds (Fig. 6C). Of these 6, only three identified the same interaction interface (RNH-1.3, PIRE-2, and PIRE-6). Of the previously known and experimentally supported physical interactors, three were identified by AF 2 but not by AF 3 (MUT-16, PIK-1, and RDE-8). Two others (MUT-7 and PRG-1) could not be identified by AF 2 even after five different runs (i.e, among 125 models), suggesting that these are either indirect interactors or require additional multimerization for complex formation. The extensive regulation of RDE-3 suggested by these predicted interactions is consistent with recent experimental results that have revealed differences in the patterns of poly-UG RNAs detected when a germline (45) or somatic gene (20) is targeted by dsRNA, and the diversity of poly-UG patterns associated with different forms of heritable RNA silencing (56).

**Figure 6.**
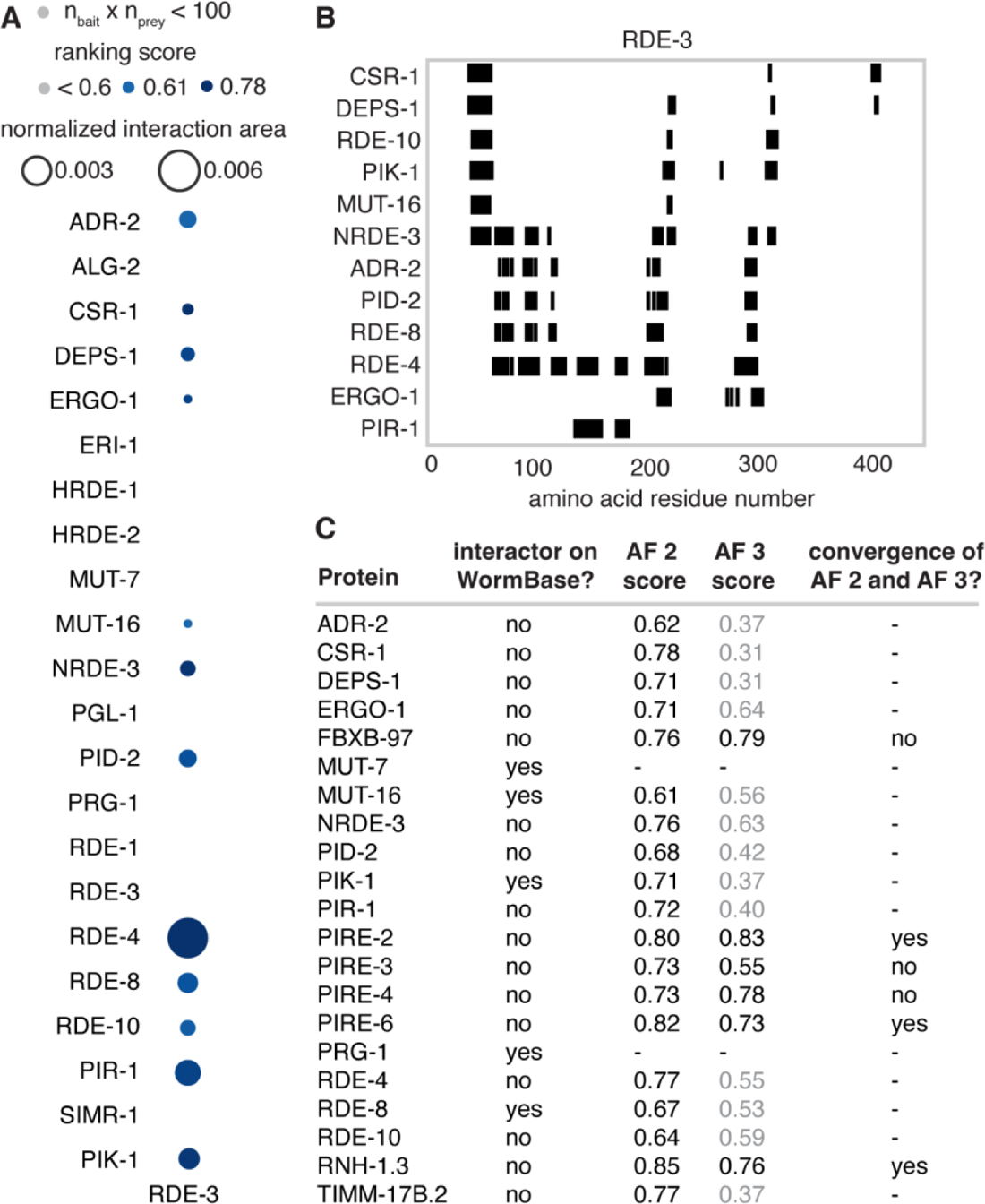
The poly-UG polymerase RDE-3 is predicted to interact with multiple proteins. (*A*) Predicted interactions of RDE-3 with known regulators of RNA silencing and the 5 proteins listed as physical interactors on WormBase (MUT-16, MUT-7, PIK-1, RDE-8, and PRG-1) identified by AlphaFold 2.3 are shown. Sizes of circles indicate normalized interaction area and shading indicates ranking score. Grey indicates ranking scores < 0.6 and/or the products of numbers of constrained residues in RDE-3 and its interactors (n_bait_ x n_prey_) < 100. Also see Movies S61 to S72. (*B*) Regions of RDE-3 protein sequence constrained by the interacting regulators. Markers (black) are as in Fig. 3B. (*C*) Table summarizing interactors of RDE-3. Experimentally identified physical interactors (interactor on WormBase?), highest score of AF 2 predicted interactions that are > 0.6 (25 models from 1 run), highest score among AF 3 predicted interactions (25 models from 5 runs), and whether the AF 2 and AF 3 structures are similar (convergence of AF 2 and AF 3?) are indicated. Scores of AF 3 models that lack any interactions between the two proteins with a predicted aligned error < 5Å and a distance < 6Å are indicated in grey.

### Predictions after an immunoprecipitation could identify direct links to other processes

Interactions identified using immunoprecipitation followed by mass spectrometry are likely to be strong interactions with abundant proteins but can be direct or indirect. Immunoprecipitation experiments can also result in large lists of putative interacting proteins (e.g., 365 for CSR-1 in one study (54)), which can make it challenging to prioritize the interactors for further study. Examining candidate interactors using AlphaFold is potentially a way to distinguish direct interactors from indirect interactors or spurious co-precipitates.

To test this possibility, we chose a relatively selective immunoprecipitation experiment that identified 12 putative interactors of the Z-granule surface protein PID-2 (36). Of this dozen, 11 were ∼1000 aa or smaller and therefore amenable to testing using AF 2 with reasonable computational resources. Five were identified as significant interactors with the following ranking scores: PID-5 – 0.63, PID-4 – 0.63, KIN-19 – 0.74, PAR-5 – 0.74, and T07C4.3 – 0.53 (Fig. S5*A*; Movies S73 to S77). Even considering only the 4 proteins that satisfy the more stringent criterion of >0.6 ranking score, this analysis provides useful information. First, it identifies PID-3 and PID-4 as direct interactors in agreement with further experimental evidence provided in the study (36) and predicts the sets of residues constrained by the interactions (Fig. S5*B*), which can be tested using additional experiments. Second, it suggests that KIN-19 and PAR-5 are additional direct interactors. KIN-19 is an ortholog of Casein kinase and was recently shown to phosphorylate the Argonaute ALG-1 (57). The predicted interaction with PID-2 suggests a wider role for this kinase in the regulation of RNA silencing, potentially through the phosphorylation of PID-2 or other substrates localized near Z granules. PAR-5 is a 14-3-3 protein required for the proper partitioning of cytoplasmic components in the early embryo (58). Furthermore, PAR-5 does not show a significant interaction with 19 other tested regulators of RNA silencing after one AF 2 run (Fig. 7A), is frequently identified as interacting with PID-2 by AF 2 (Fig. 7B), and has an extensive interaction interface (Fig. 7C) that constrains the C-terminal 17 amino acids of PID-2 (Fig. 7D). Underscoring the high confidence in this interaction, the same interaction is also predicted by AF 3 (Fig. 7E and Movie S78) and the **R**GF**S**_450_EC**P** sequence within the interaction domain is close to a consensus (RxxpSxP) for binding 14-3-3 domains (59) with S_450_ phosphorylated by an atypical Protein Kinase C (60). Nevertheless, determining if, when, and where any predicted interactions occur in vivo will require many future experiments.

**Figure 7.**
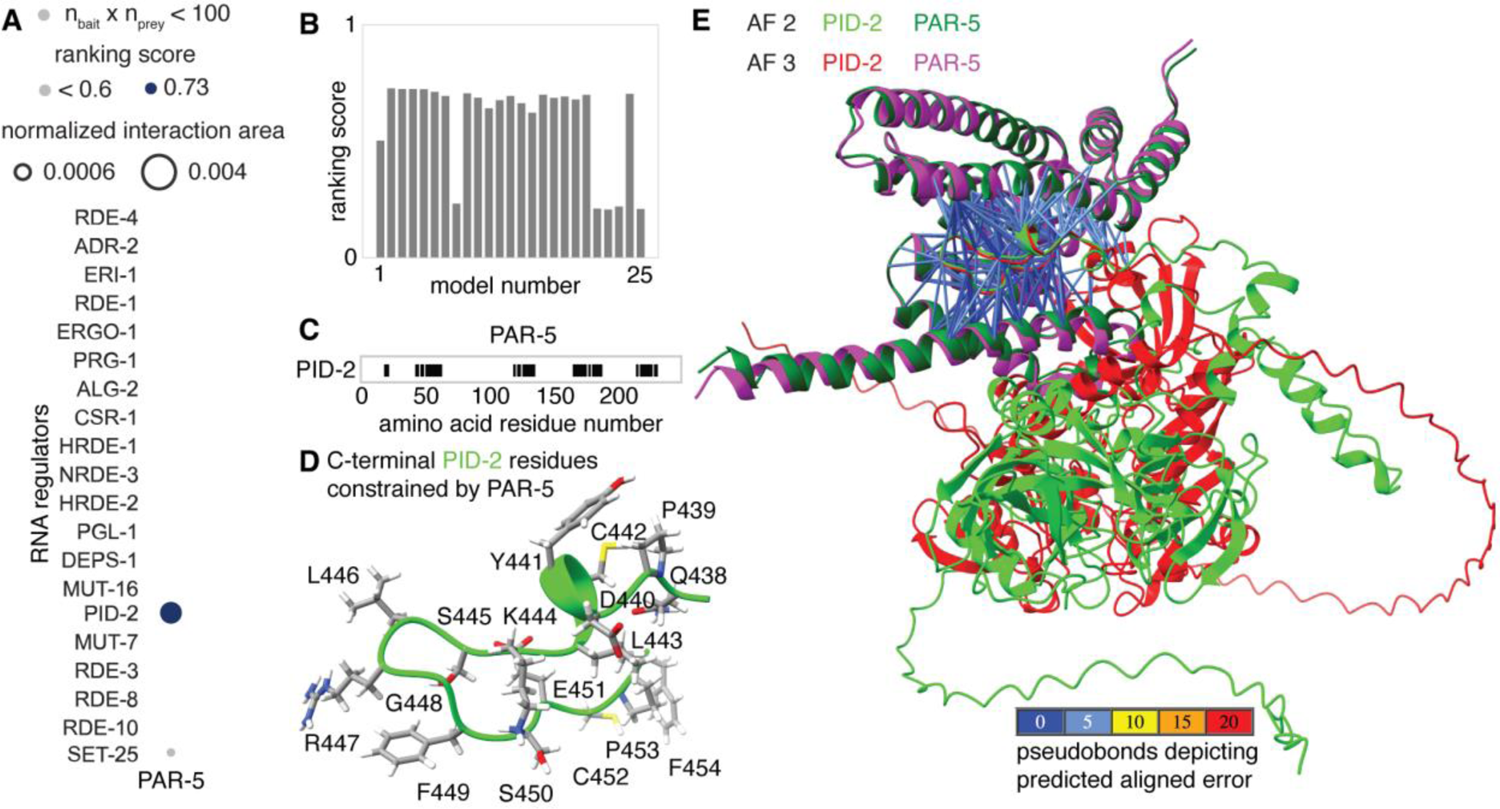
PAR-5 is predicted to interact with the Z-granule surface protein PID-2/ZSP-1 but not with many other tested regulators of RNA silencing. (*A*) Predicted interactions of PAR-5 with known regulators of RNA silencing identified by AlphaFold 2.3 are shown. Area of circles and shading are as in Fig. 6A. (*B*) Distribution of ranking scores for the 25 models of PAR-5:PID-2 generated by AF 2. (*C*) Regions of PAR-5 protein sequence constrained by interactions with PID-2. Markers (black) are as in Fig. 3B. (*D*) Structure of the C-terminus of PID-2 constrained by PAR-5. (*E*) Overlay of models predicted by AF 2 and AF 3 superimposed using PAR-5 showing similar interactions between the C-terminus of PID-2 (lime or red) and PAR-5 (magenta or green) although the rest of the PID-2 protein are positioned differently in the two models. Pseudobonds are as in Fig. 3A. Also see Movie S78.

### Some predicted interactions could be challenging to demonstrate experimentally

Obtaining experimental support for direct interactions between proteins can be difficult. For example, an interaction between the most abundant Gα protein in the brain (GαχσΣυβΣταρτοχσΣυβενδ (61), GOA-1 in *C. elegans*) and the diacylglycerol kinase DGK-1 is strongly predicted by genetic analysis (62,63). Both AF 2 and AF 3 predict the same extensive binding between GOA-1 and DGK-1 (Fig. S6 and Movie S79). Furthermore, the interaction interface is largely preserved and reliably predicted by AlphaFold 3 when GOA-1 is by itself or bound to either GTP or GDP (Fig. S6 and Movie S79). Yet, early attempts using purified proteins failed to reveal a detectable interaction between DGK-1 and GOA-1 in vitro (64), and this interaction has remained a conjecture for more than two decades.

While biochemical approaches rely on preserving or recreating in vitro the unknown conditions in vivo to coax a detectable interaction between proteins, prediction algorithms that incorporate extensive multiple sequence alignments (e.g., AF 2 and to an unknown extent AF 3) can use the co-evolution of residues to deduce the interaction. Given these complementary strengths, systematic analyses using both multiple experimental approaches (1) and multiple prediction algorithms are needed to find the edge of predictability for protein-protein interactions.

### The top 100 genes include many that could link RNA silencing to other processes

To examine if the observations above would hold when analyzing a larger set of genes, we examined the top 100 genes ordered according to their *r*_100_ values. To quantify the correlated presence or absence of genes in different lists we used a measure of mutual information (7) named here as historical mutual information (HMI) to emphasize the subjective nature of this measure because it depends on both functional relatedness of the genes and biased availability or inclusion of data (see supplementary methods and the 6_HMI_explorer.py program for exploring clusters of genes interactively). Using HMI to cluster these genes revealed three major clusters (64, 20, and 14 genes) and two other unconnected genes (Fig. 8A, Table S2) when communities are formed with a threshold distance (1-HMI) of 0.9 or less for a link between two genes.

**Figure 8.**
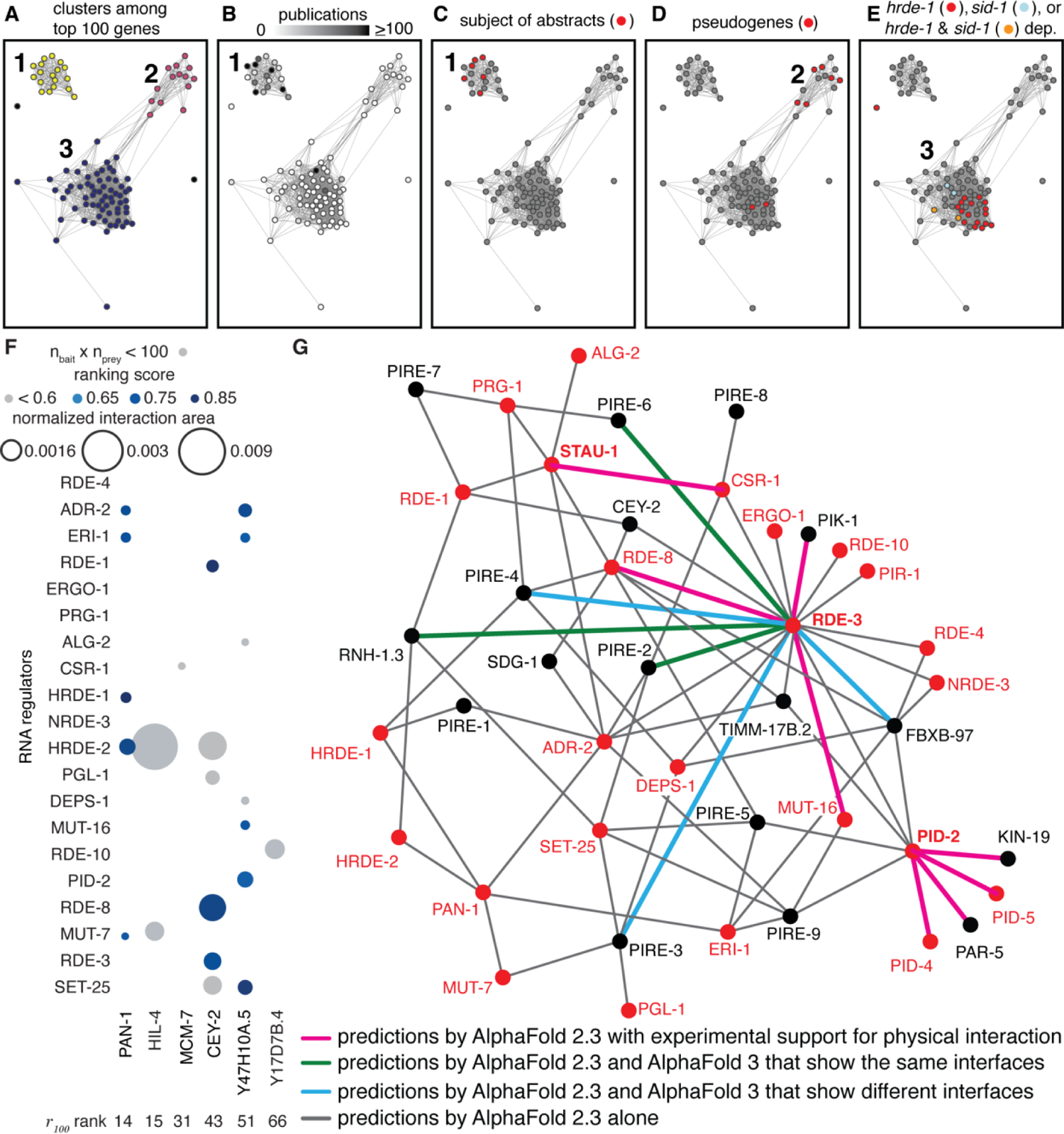
Clusters formed by understudied regulated genes suggest priorities for detailed study. (*A* to *E*) Properties of the top 100 regulated genes in the field of RNA silencing in *C. elegans.* (*A*) Clusters of genes based on their historical mutual information (HMI). Threshold for link: distance (1 - HMI) < 0.9. (*B* to *E*) Network in (*A*) with nodes colored to show number of publications per gene (white, 0; black, ≥100) (*B*), genes that have been the main subject of abstracts on RNA silencing in *C. elegans* (*C*), pseudogenes (red) (*D*), and genes changed in *hrde-1* mutants (69) (red), a *sid-1* mutant (16) (cyan), or both (orange) (*E*). (*F*) Predicted interactions of proteins encoded by genes with different *r_100_* ranks with known regulators of RNA silencing. Sizes of circles indicate normalized interaction area and shading indicates ranking score. Grey indicates ranking scores < 0.6 and/or the products of numbers of constrained residues (n_bait_ x n_prey_) < 100. Also see Movies S80 to Movie S93. (*G*) All interactions (connecting lines) depicted were identified by AF 2 (grey). Some are supported by experimental evidence for physical interaction (magenta) and some are also predicted by AF 3 with either similar (green) or different (cyan) interfaces. Known regulators of RNA silencing are in red and those used as baits to look for predicted interactors (STAU-1, PID-2, and RDE-3) are in bold. Also see Table S4.

Only one cluster (cluster 1 in Fig. 8A) had significant numbers of genes associated with gene ontology terms (Table S3). These genes encode proteins involved in RNA silencing and/or play roles in other processes such as cell division and germ cell development. Consistently, this cluster also had the greatest number of genes that have been described in multiple publications (Fig. 8B), including all the genes that have been featured in abstracts on RNA silencing (Fig. 8C). Therefore, the analysis of additional genes in this cluster could be relevant for RNA silencing and connect it to other processes (e.g., the cell cycle). Several predicted interactions are consistent with this speculation. One, among the other genes in cluster 1 is the gene encoding PAR-5, which is predicted to selectively interact with the known regulator of RNA silencing PID-2 (Fig. 7). The predicted interaction could have a role in segregating PID-2, and potentially other components of Z-granules, to the posterior side before the first cell division during embryonic development (58). Two, cluster 1 also includes the gene encoding MCM-7, which is predicted to selectively interact with CSR-1 (ranking score 0.59 in Fig. 8F; also see Fig. S7 and Movie S80). This interaction is also supported by an immunoprecipitation experiment (54) and it could have a role in the chromosome segregation function of CSR-1 (65) because of the established role of the MCM complex in DNA replication (66). Three, two other proteins encoded by genes in this cluster that were also tested (CEY-2 and PAN-1) are predicted to interact with some RNA regulators (8 of 40 potential interactions tested in Fig. 8F; also see Fig. S7 and Movies S81 to S88).

Since six of the eight pseudogenes are in a small cluster (Fig. 8E, 6 of 14 genes in cluster 2), the other genes in this cluster could potentially be targets of regulation without specific downstream regulation or be co-regulated sensors of pseudogene RNA levels. Two PIRE proteins (PIRE-3/E01G4.5 and PIRE-5/K02E2.6) are also present in this cluster. Another gene Y47H10A.5 encodes a protein with similarity to decapping nuclease (Foldseek, E-value < 0.05) and is targeted by miR-243 (67), leading to RDE-1-dependent small RNA production (67,68). Since, the Y47H10A.5 protein is predicted to interact with multiple regulators of RNA silencing (Fig. 8F, Table S2, Fig. S7, and Movies S89 to S93), we refer to it as PIRE-9.

There is a large overlap between a set of genes that require HRDE-1 for downregulation (67 genes in both replicates from worms grown at 15°C (69)) and genes in a single cluster (Fig. 8F, 20 of 64 genes in cluster 3). One possible explanation for this abundance and clustering could be that *hrde-1*-dependent gene lists are among the most numerous generated by the field and/or included in our analysis (42 of 398 lists). Alternatively, genes that are subject to HRDE-1-dependent silencing could be extensively regulated by many other regulators and require this additional downregulation for fitness – i.e., overexpression of these genes is detrimental. Consistent with this possibility, loss of HRDE-1 results in progressive sterility that can be reversed by restoring HRDE-1 activity (69). Also, as expected for the use of HRDE-1 downstream of SID-1, genes upregulated using *sid-1* (18 genes in animals lacking *sid-1* (16)) overlap with genes in the same cluster (Fig. 8F and Table S2, 4 of 64 in cluster 3).

In addition to these hypotheses, how interacting with PIRE proteins modulates the functions of known regulators of RNA silencing could be experimentally tested (Table S3). Future studies by labs working on multiple aspects of RNA silencing in *C. elegans* have the potential to test and enrich the classification of the regulated yet understudied genes revealed here, including by identifying many more PIRE proteins.

## Discussion

Our analysis has identified selectively regulated yet understudied genes in the field of RNA silencing in *C. elegans*, some of which encode predicted influencers of RNA-regulated expression that act through protein-protein interactions. To facilitate easy inspection of all the predicted interactions identified in this study, we generated a network diagram (Fig. 8G) that summarizes the 77 predicted interactions among 42 interactors with varying amounts of support. Minimally, all interactions shown are predicted by AlphaFold 2.3. A survey of the information available on WormBase for all 42 interactors revealed that 8 of these interactions are already supported by some experimental evidence for physical interaction. We note that this is not an exhaustive list of all possible interactions even among the 42 interactors considered and expect that future experimental work will refine this view.

### The inevitable bias of progress

Bias during progress in a field is unavoidable and its causes are complex, including availability of technology, researcher pre-disposition, perceived importance of a direction, current societal need, etc. Therefore, the comprehensive appraisal of a field through equal representation of all important aspects is impractical. Indeed, our analysis involved the manual collation of many datasets for comparison, which could have resulted in omissions and inclusions that spark disagreements. While future extensions of this work could automate the process of aggregating and comparing data, flexible inclusion of different lists in the analysis would be needed to enable customization based on the expertise, interests, and risk tolerance of individual labs. Furthermore, earlier studies using older technologies could have led to conclusions that need revision. For example, when analyzed using multi-copy transgenes, the dsRNA-binding protein RDE-4 showed a cell non-autonomous effect (70,71), but when analyzed using single-copy transgenes, RDE-4 showed a cell autonomous effect (72). Since different researchers could interpret such conflicting data differently (e.g., differences in levels of tissue-restricted expression versus differences in extent of misexpression in other tissues), it is useful to preserve the ability to customize lists. With the expanding number of lists generated by large-scale experimental approaches in different fields, identifying selectively regulated yet understudied genes could aid the prioritization of genes for detailed mechanistic studies using the limited resources and time available for any lab.

### Function(s) of the *x*-dependent gene

Different properties of a single protein or RNA could be important for different biological roles (73,74), or the same properties could be important for different processes. Despite such variety, a gene found in many lists could become associated with a single label because of the historical sequence of discovery (e.g., HRDE-1-dependent genes; many in cluster 3, Fig. 8E), thereby obscuring additional roles of that gene. Most of the PIRE proteins are predicted to interact with more than one tested regulator of RNA silencing (e.g., ten in Fig. S3). If these interactions are validated through experimental analyses, it will not be possible to classify these PIRE proteins into single pathways. Indeed it can be challenging to delineate pathways when multiple regulators in an intersecting network make quantitative contributions to an observed effect (20). The well-recognized difficulty in defining the function of a gene (75) is exacerbated in these cases, making it more appropriate to consider these proteins as entities within a system whose roles depend on context (see (76) for similar ideas).

### Metrics for historically contingent progress

Exhaustive collation of past progress can be difficult because of the many formats in which data and inferences are presented. Of these, tabular data are particularly amenable to future computation. The simple *r_g_* metric provides a weighted sum of frequently occurring features (e.g., genes) for prioritizing the top 25 genes (Fig. 1) or 100 genes (Fig. 8). However, the number of genes considered for calculating *r_g_* can influence the prioritized set obtained (Table S4). Specifically, the same genes were identified as the top 25 genes by considering 1000 genes (1000^th^ *r_1000_* gene being present in 53 lists) or 100 genes (100^th^ *r_100_* gene being present in 72 lists), and 16 of these genes were identified by considering 25 genes (25^th^ *r_25_* gene being present in 84 lists). More complicated metrics that consider other useful aspects of the data such as effect size (77) of the reported change (e.g., measured for fold-change when using RNA-seq), discoverability of the change using a technique (e.g., influenced by abundance of a protein for immunoprecipitation), and reliability of the technique used (e.g., adequacy of replicates for estimating noise) could be developed in the future to extract more information. Historical mutual information provides a measure of predictability that is an unknown mix of functional relatedness and biased attention, hence ‘historical’. This metric is simply a normalized measure of mutual information (78), which captures the predictability of one feature given knowledge about another feature and is widely used (79) because of the ability to capture both correlations and anticorrelations without any knowledge of underlying causality. The tendency to progress by building upon past discoveries and to communicate by connecting to concepts of perceived importance makes the growth of knowledge akin to growth of networks through preferential attachment (see simulation in Movie S94). Metrics that take advantage of this aspect could be developed to reduce bias in the information (e.g., by weighting based on community size). However, separating features that appear to be important based on progress in a field from what is inherently important given the characteristics of a system can be challenging.

### From transcript changes to protein-protein interactions

Positive feedback loops that drive growth and development are a ubiquitous feature of life (80). Yet, living systems are also characterized by homeostasis (4), which needs negative feedback to suppress runaway processes. For example, in a chain of biochemical reactions, product inhibition (81) can be used to regulate production to match need. While this organization enables compensation in response to change, complete compensation for all processes is clearly not possible as evidenced by the fact that many mutations have measurable consequences. A specific case of this general principle is transcriptional adaptation, where the mutation-induced degradation of a transcript results in compensatory changes in the levels of other transcripts (82). The existence of PIRE proteins suggests that another way for organisms to compensate for the perturbation of a protein that regulates a process is to change the levels of other proteins that can regulate the same process through protein-protein interactions. Thus, we speculate that perturbing a protein could sometimes alter the mRNA levels of its interactors because of the prevalence of feedback regulation in living systems. If true, this feature of life provides a strategy for combining RNA sequencing and protein structure predictions to identify protein-protein interactions of regulatory importance.

## Data Availability

All scripts used in this study are available at GitHub (AntonyJose-Lab/Lalit_Jose_2024) and has been archived at Zenodo (https://zenodo.org/records/13952718).

## Acknowledgements

We thank Tom Kocher, Brian Pierce, and members of the Jose lab for comments on the manuscript; four anonymous reviewers for their feedback and suggestions; Brian Pierce for discussions and Rui Yin for installing AlphaFold 2 on the UMD HPCC; and Carlos Retamal and José Feijo for getting us started with AlphaFold. We acknowledge the University of Maryland supercomputing resources (http://hpcc.umd.edu) made available for conducting the research reported in this paper.

