## Supplemental Information for "Selecting genes for analysis using historically contingent progress: from RNA changes to protein-protein interactions"

**Affiliations:**

**This file includes:**

Supplementary Methods
SI References
7 Supplementary Figures
5 Supplementary Tables
94 Supplementary Movie Legends

### Supplementary Methods

**Analysis of gene data tables.** To identify studies on RNA silencing in *C. elegans* with data tables that can be compared across all studies, we used the term 'C. elegans RNA silencing' to search PubMed. After examining the abstracts of more than 2000 studies that resulted from the search, the available data tables from 82 studies that were published between 2007 and 2022 were downloaded (Table S1), reformatted into 398 distinct tables manually and/or using custom scripts. Metadata if supplied by the authors for each table were retained as comments above each table. Gene names were unified using the Gene Name Sanitizer (https://wormbase.org/tools/mine/gene_sanitizer.cgi) as on 26 April 2022 (0_sanitization.py). It is unclear how an exhaustive list of papers that is nevertheless field-restricted could ever be defined for any field. Accordingly, our list of RNA silencing studies in *C. elegans* is not exhaustive and we apologize to colleagues whose work is not included in our analysis. Nevertheless, this effort captured additional datasets compared with those available in other more unrestricted collections that attempt to collect tables from all studies on an organism (e.g. WormExp 2.0 (1)). Only 30 of the 55 studies published before 2017 and included in this study overlapped with the 461 included in WormExp 2.0 as on 27 Jul 2017, which was available for download from the website (https://wormexp.zoologie.uni-kiel.de/wormexp/). This overlap was determined by comparing the paper IDs using a custom script (0_dataset_wormexp_overlap.ipynb). Data tables that reported p-values or adjusted p-values were filtered to only include entries with p < 0.05 (0_filter_pvals.py). Since fold-changes were not always available, for every dataset, genes were scored as present or absent (1_TableOccupancy.py) to generate a heatmap featuring the most frequently changed genes sorted by $r_g$ values, where the number of genes considered ($g$) can be arbitrary (e.g., 25 in Fig. 1*F* and 100 in Fig. 8). The relationships between the parameters $S_i$, $T_i$, and $g$ (Fig. 1*C*) were obtained using simulated data by sampling 100 random sets of genes (0_rg_simulation_box-whisker.ipynb) as the top $g$ genes from a total of 20,000 genes and similarly sampling the genes in datasets of various sizes ($T_i$). For each gene in published lists in the field, the number of references listed on Wormbase (https://wormbase.org/) was used as a measure of the extent to which the gene has been studied (2_fig1D_r25_references.ipynb). Genes with fewer than 10 references were defined as understudied (Fig. 1*D*). To generate the heatmap (3_fig1F_fig1G_r25_related.ipynb, 4_heatmaps_normalized_100_full.ipynb), genes were ordered by decreasing values of $r_{25}$ (top to bottom in Fig. 1*F*) and datasets were ordered by decreasing values of $\frac{S_i}{T_i}$. (left to right in Fig. 1*F*). To determine the co-occurrence patterns of all pairs of genes, Jaccard distances ($d_J = 1 - \frac{|X \cap Y|}{|X \cup Y|}$, where $X$ and $Y$ are sets of lists containing genes $x$ and $y$, respectively) were calculated for each pair and all genes were hierarchically clustered using the 'average' linkage method. Relationships between genes based on occurrence in datasets were also captured as normalized mutual information (5_sklearn_nmi.ipynb) and defined as historical mutual information (HMI) to emphasize the dependence on the biased availability or inclusion of data based on historical progress in addition to the functional relatedness of the genes. Specifically, it was defined to be a symmetric and normalized mutual information score (2) and was calculated using the function normalized_mutual_info_score from scikit-learn (3) for genes $X$ and $Y$:

$$HMI(X;Y) := \frac{2.MI(X;Y)}{H(X) + H(Y)},$$

where $MI(X;Y) = \sum_y \sum_x P_{(X,Y)}(x,y)\, log_2\left(\frac{P_{(X,Y)}(x,y)}{P_X(x)P_Y(y)}\right)$, $H(X) = -\sum_x P(x) log_2\big(P(x)\big)$, and $H(Y) = -\sum_y P(y) log_2\big(P(y)\big)$. Mutual information (MI) determines how different the joint distribution of the gene pair (X, Y) is from the product of the marginal distributions of each gene, $H(X)$ and $H(Y)$ are the entropies of the two genes, and $P(\ldots)$ indicates probabilities. Clusters of genes based on HMI values were identified using the Girvan-Newman algorithm (4). An interactive graphical user interface (GUI) for visualizing clusters and genes of interest (6_HMI_explorer.py) was created using Dash (Python) and figures highlighting genes within the clusters were generated (7_fig8.ipynb). Gene Ontology (GO) analysis was performed on all clusters using the Gene Ontology Resource ((5,6); https://geneontology.org/). Tables of the top 25 genes ranked by $r_{25}$ when different numbers of total top genes are considered (Table S5) were generated for comparison (8_table_S5_r25_with_100_total.ipynb, 8_table_S5_r25_with_1000_total.ipynb).

**Analysis of predicted protein structures.** Predicted protein-protein interactions were examined using Alphafold 2.3.2 and the Alphafold 3 server, downloaded to a local machine, and analyzed using ChimeraX and custom scripts.

*Alphafold 2.* For each understudied regulated gene, files with protein sequences (.fasta) encoded by the longest transcript isoform were obtained from Wormbase and combined using the program 'fasta_assembly_for_alphafold_dimer.py' to create paired fasta files to be used for testing the potential for an interaction between the two proteins. Batches of potential interactors prepared in this way were run on the high-performance computing cluster (Zaratan, UMD) using a batch submission script ('alphafold_multimer_batch_submission.sh') that modifies another script for submitting alphafold 2.3.2 jobs with the model_preset flag set to 'multimer' ('alphafold_multimer.sh'). Typical resource requests included a wall time of 18 hours, one A100 GPU, and 8 CPUs at 6 GB each. Upon completion, a script for reducing the results folder to keep only the highest-ranking model was run ('alphafold_results_cleanup.sh') before downloading from the HPCC to a local machine. To analyze and annotate the downloaded models, the 'alphafold2_dimer_batch_computed_on_zaratan.py' program and run using the command 'chimerax --exit alphafold2_dimer_batch_computed_on_zaratan.py', which runs the python program within ChimeraX-1.7.1. Data for all predicted interactions to be analyzed together were collected under the same file ('yyyy_m_d_alphafold2_summary_stats'), where yyyy_m_d indicates date. This program also generated most of the supplemental movies. The program 'predicted_influencer_of_RNA_regulated_expression_d2.py' was then run to extract information about the interactions and make plots with either absolute interaction areas or areas normalized based on the sizes of the interacting proteins (passed to the program through the files ('yyyy_m_d_A_list_sizes' and 'yyyy_m_d_B_list_sizes'). Additional plots showing the residue numbers and locations of residues interacting with each regulator were created using the program 'interactor_map_for_a_protein_with_another_set_of_proteins.py'. The final figure showing the scaled area of interaction shaded according to the ranking score (Fig. 2*B*) was generated using 'final_interactors_filtered_by_model_rankings.py'.

Analysis of Alphafold 2 models using chimeraX and the downstream computations can also be performed using the two scripts 'predicted_dimer_chimerax.py' and 'predicted_dimer_python.py'. These streamlined scripts also generate distributions of the ranking scores for the 25 models identified with each run of Alphafold 2.

*Alphafold 3*. Essentially the same workflow as above was used after downloading the predicted interactions for pairs of proteins from the Alphafold 3 server, which was run in batches of 10 or 20 per day based on quota availability. Parsing the resulting data required some minor modifications to the programs because the error files (.json) and the structure files (.cif) were in different formats and labeled differently. The program 'alphafold3_dimer_batch_computed_on_google.py' was used for analyzing these predictions.

*Comparisons of Alphafold 2 and Alphafold 3*. For comparisons of the two prediction approaches, the 'alphafold3_dimer_batch_computed_on_google_comparing_af2_af3.py', 'predicted_influencer_of_RNA_regulated_expression_d2_af2_vs_af3_af3_run.py' and 'interactor_map_for_a_protein_with_another_set_of_proteins_comparing_af2_vs_af3_rerun_on_af3.py' programs were used.

*Illustrations.* Illustrations of protein-protein complexes for figures were created manually using ChimeraX (1.7.1 or 1.8-rc2024.05.24) and Adobe Illustrator (28.5). Typical workflow on ChimeraX included opening the .pdb or .cif files and the associated predicted aligned error files (.json or .pkl), aligning them as necessary, coloring different proteins, overlaying multiple models when relevant, and adding inter-protein pseudobonds based on criteria before saving images and/or movies. All interactions predicted in the study were summarized using Gephi (v. 0.10.1 202301172018) and Adobe Illustrator (v. 28.7.1).

### Supplementary Figures

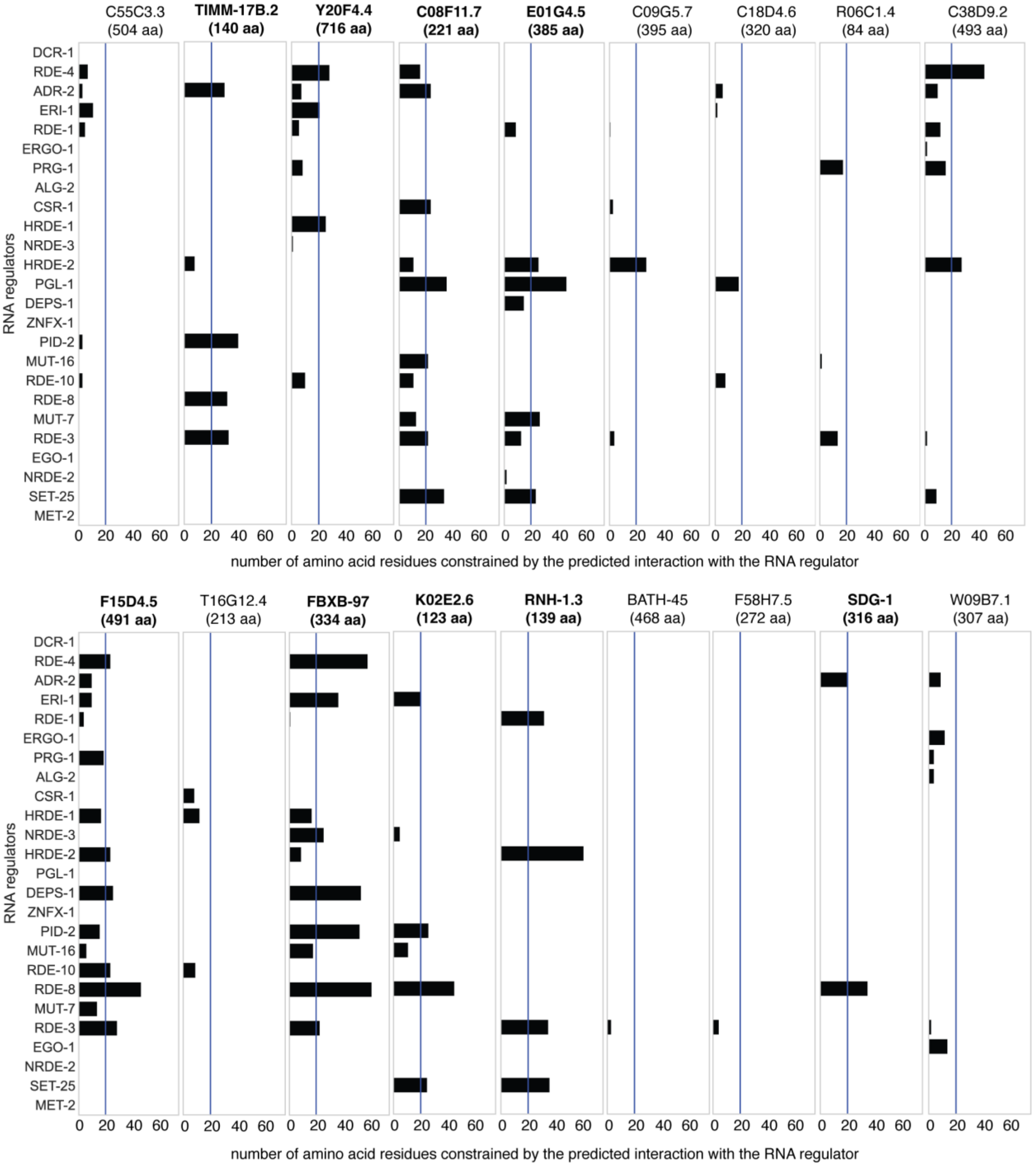


**Figure S1. Numbers of candidate PIRE protein residues constrained by the predicted interacting regulator of RNA silencing in *C. elegans*.** Numbers of residues that interact with an inter-protein PAE < 5Å and a distance between residues < 6Å are plotted for each interaction between a protein encoded by an understudied gene and a known regulator of RNA silencing in *C. elegans*. A threshold of 20 residues (blue line) and a ranking score >0.6 was used to separate candidate PIRE proteins (highlighted in bold) from others encoded by understudied genes.

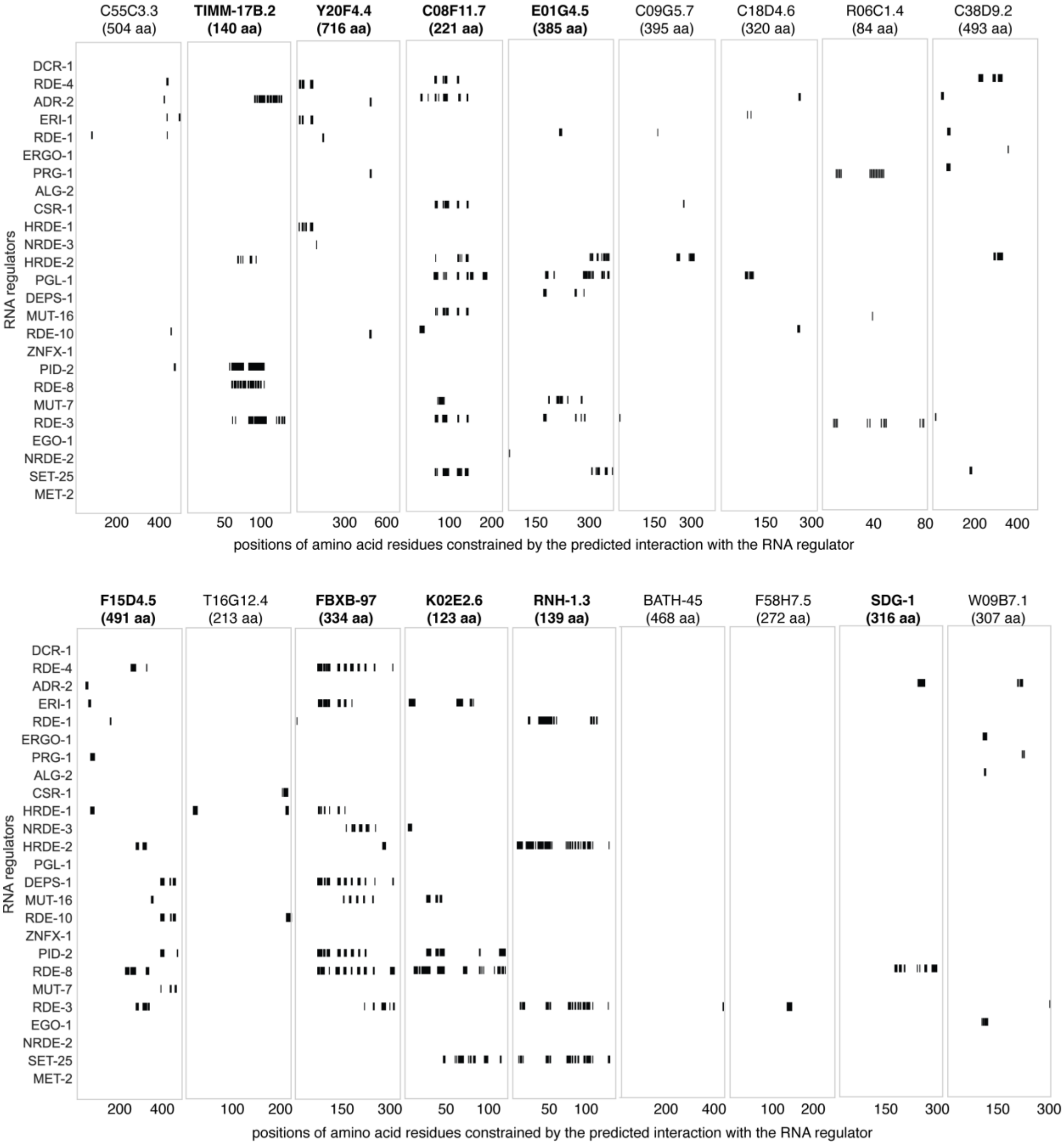


**Figure S2. Regions of the candidate PIRE protein sequence constrained by the predicted interacting regulator of RNA silencing in *C. elegans*.** Markers (black) are enlarged with respect to the X-axis for visibility (e.g., the marker denoting the interaction between RDE-1 and FBXB-97 only indicates one residue). Understudied genes that encode candidate PIRE proteins are highlighted in bold.

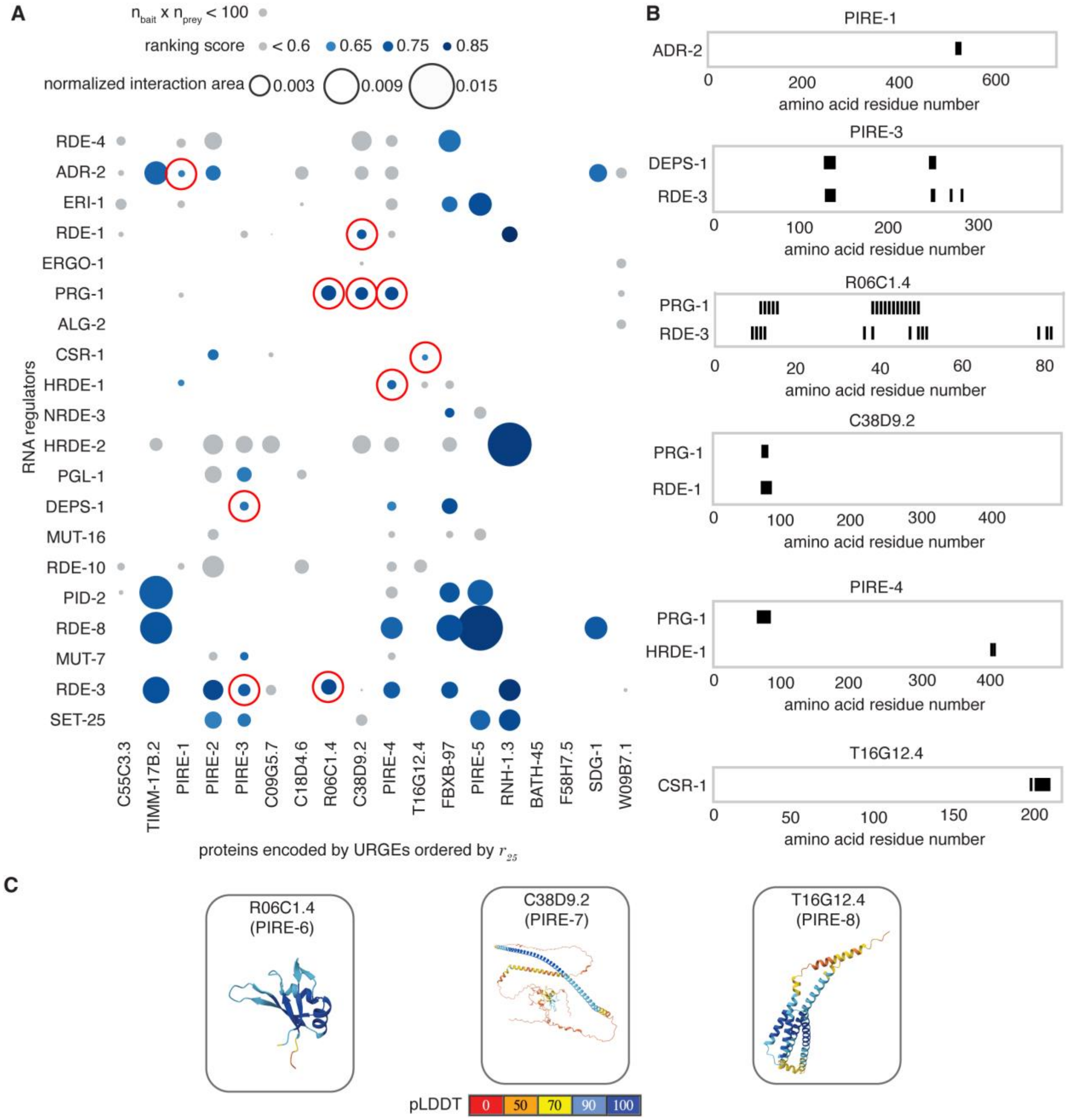


**Figure S3. Predicted interactions between proteins encoded by the top 25 genes and known regulators of RNA silencing identified with more permissive criteria.** (*A*) The area of the interaction surface between partners normalized by the product of the sizes of the interactors is shown as a bubble plot. Interactions are shaded according to ranking score. Interactions for which the product of the numbers of interacting residues ($n_{bait}$ x $n_{prey}$) with an inter-protein predicted aligned error < 5Å and inter-residue distance < 6Å in a model with a ranking score > 0.6 is less than 100 are shaded grey. Ten interactions identified in addition to those found using criteria in Fig. 2*B* are highlighted with red circles. (*B*) Regions of the additionally identified proteins constrained by the interacting regulators (red circles in *A*) with markers depicted as in Fig. 3*B*. (*C*) Predicted structures for additional PIRE proteins with pLDDT as in Fig. 2*D*. Also see Movies S39 to S48.

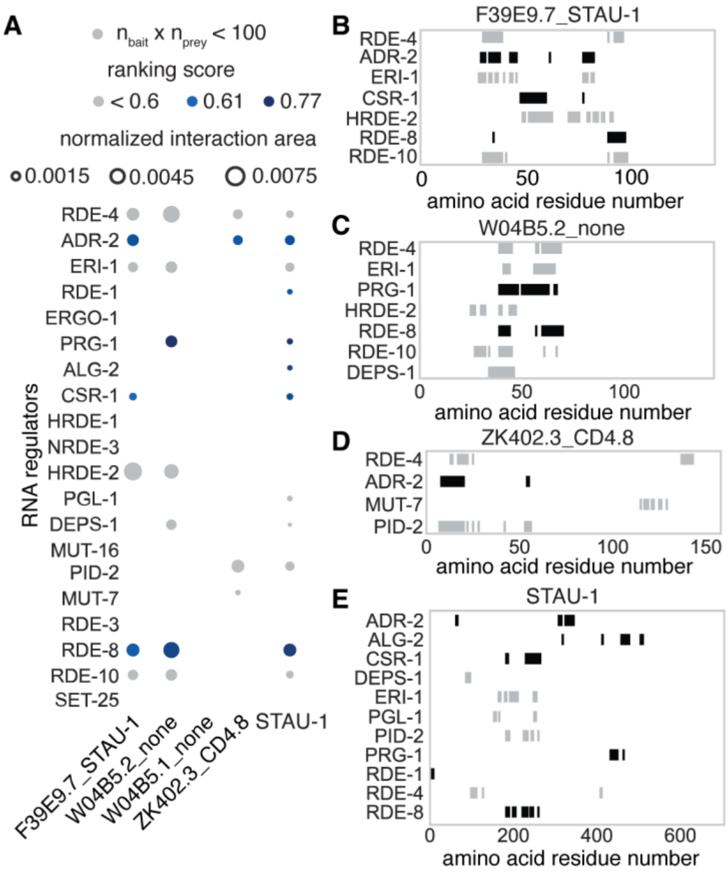


**Figure S4. Predicted interactions of potential peptides encoded by pseudogenes and a homologous protein with regulators of RNA silencing.** (*A*) Interactions are depicted as in Fig. S3*A*. The proteins are labeled with their closest BLAST matches separated by an underscore (e.g., F39E9.7_STAU-1 indicates that the peptide that could be encoded by F39E9.7 shares homology with STAU-1). (*B* to *D*) Regions of the longest peptide sequences encoded by F39E9.7 (*B*), W04B5.2 (*C*), and ZK402.3 (*D*) constrained by the interactors are shown with markers as in Fig. 3*B*. (*E*) Regions of STAU-1 protein sequence constrained by interacting regulators of RNA silencing are shown with markers as in Fig. 3*B*. Also see Movies S49 to S60.

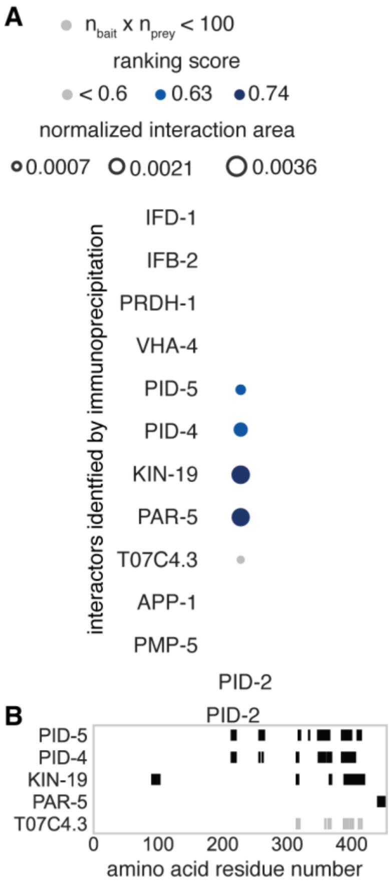


**Figure S5. Predicted interactions between PID-2/ZSP-1 and proteins identified in a pulldown of PID-2 (as reported in (7)).** (*A*) Interactions are depicted as in Fig. S3*A*. (*B*) Regions of the PID-2 protein sequence constrained by the interactors are shown with markers as in Fig. 3*B*. Also see Movies S73 to S77.

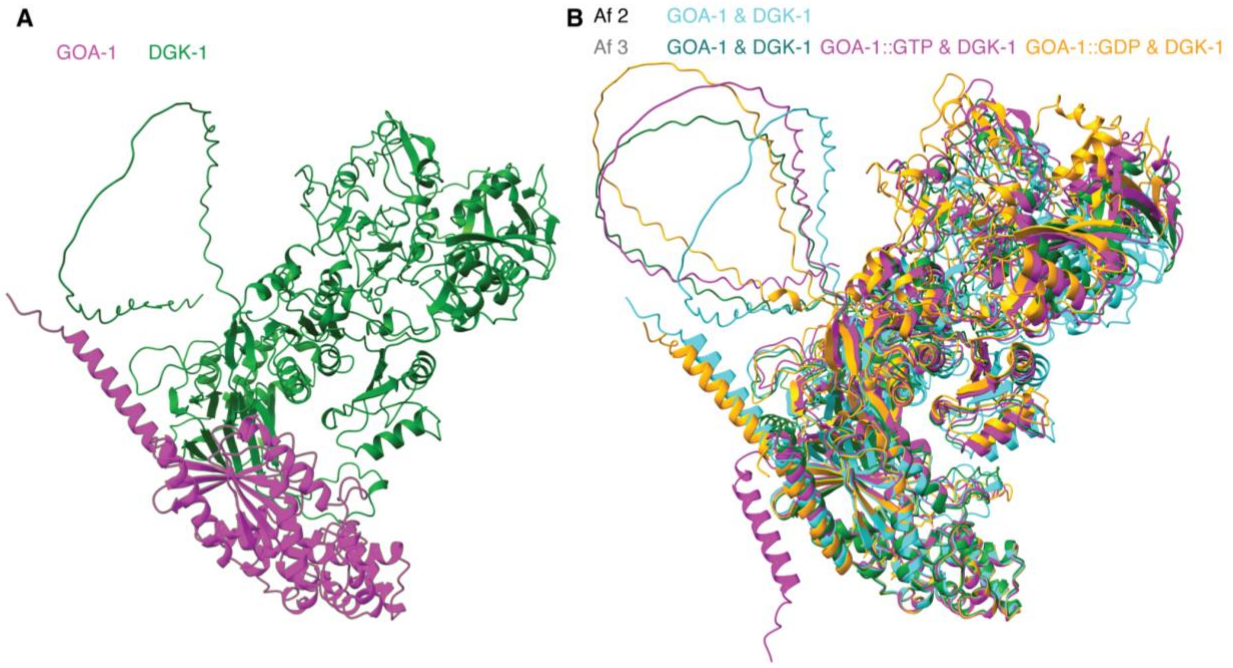

**Figure S6. Interactions between the G alpha protein GOA-1 and the diacylglycerol kinase DGK-1 predicted by AlphaFold.** (*A*) Interaction between GOA-1 (magenta) and DGK-1 (green) predicted by AlphaFold 2. (*B*) Overlay of the GOA-1::DGK-1 complex predicted by AlphaFold 2 (cyan) with those predicted by the AlphaFold 3 server (green, magenta, and orange for free, GTP-bound, and GDP-bound GOA-1, respectively). Also see Movie S79.

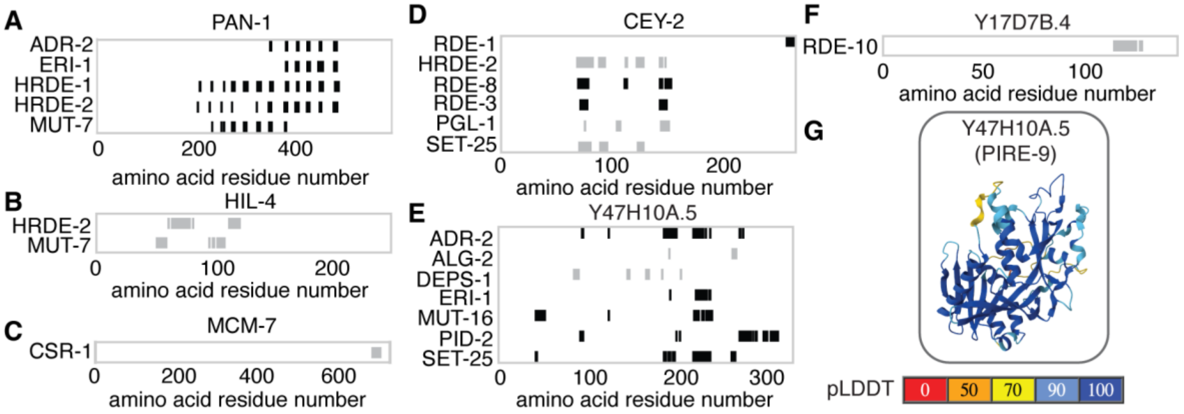


**Figure S7. A sampling of predicted interactions between regulators of RNA silencing and proteins encoded by the top 100 genes with varying $r_{100}$ ranks.** (*A* to *F*) Regions of the PAN-1 (*A*), HIL-4 (*B*), MCM-7 (*C*), CEY-2 (*D*), Y47H10A.5 (*E*), and Y17D7B.4 (*F*) protein sequences constrained by the interactors are shown with markers as in Fig. 3B. Also see Movies S80 to S93. (*G*) Predicted structures for an additional PIRE protein with pLDDT as in Fig. 2*D*. Also see Fig. 8*F*.

### Tables and Table Legends

**Table S1. Data tables used in this study.** List of the 398 tables used along with links to the 82 studies from which they were taken and a brief description of the data types. See excel file.

**Table S2. Top 100 genes grouped according to historical mutual information (HMI).** List of genes within clusters formed by the top 100 genes with distance (1 – HMI) < 0.9. The two genes that are not part of any clusters are listed as singletons.

| **Cluster 1** | **Cluster 2** | **Cluster 3** | **Singletons** |
|---|---|---|---|
| *csr-1* | F39F10.4 | *gpx-8* | R03D7.2 |
| *tbb-2* | H09G03.1 | F40D4.13 | *pyk-1* |
| *hsp-1* | W04B5.1 | *dyf-3* | |
| *cey-2* | Y47H10A.5 | C46G7.5 | |
| *pgl-3* | Y17D7B.4 | *citk-1* | |
| *hrde-1* | F39E9.7 | Y57G11C.51 | |
| *mcm-7* | ZK402.3 | *saeg-2* | |
| *klp-15* | E01G4.5 | F09C8.2 | |
| *par-5* | W04B5.2 | *gly-13* | |
| *klp-7* | W05H12.2 | *fbxb-97* | |
| *cdk-1* | K02E2.6 | Y20F4.4 | |
| *hil-4* | Y37E11B.2 | ZK973.8 | |
| *wago-1* | Y105C5A.14 | *spch-1* | |
| *hsp-90* | F55C9.3 | Y57G7A.5 | |
| *cpg-1* | | *pan-1* | |
| *rme-2* | | W09B7.1 | |
| *wago-4* | | *elf-1* | |
| *tba-2* | | C38C3.3 | |
| | | T20F7.1 | |
| | | *fkb-8* | |
| | | K05C4.9 | |
| | | F15D4.5 | |
| | | *sea-2* | |
| | | F55B11.6 | |
| | | F41G4.7 | |
| | | T16G12.8 | |
| | | C30G12.1 | |
| | | *saeg-1* | |
| | | *rnh-1.3* | |
| | | Y53F4B.5 | |
| | | E02H9.3 | |
| | | *his-24* | |
| | | ZK909.3 | |

| |
|---|
| *vet-6* |
| C04G6.6 |
| *lin-15B* |
| *qdpr-1* |
| W09B7.2 |
| K09H9.7 |
| T02G5.4 |
| *lido-18* |
| T11F9.10 |
| *scrm-4* |
| *clp-6* |
| C08F11.7 |
| *pdfr-1* |
| F58H7.5 |
| T16G12.4 |
| C09G5.7 |
| Y48G1BM.6 |
| C18D4.6 |
| ZK795.2 |
| *ceh-20* |
| W05F2.4 |
| *bath-13* |
| *timm-17B.2* |
| *fbxa-192* |
| R03H10.6 |
| *bath-45* |
| C55C3.3 |
| R06C1.4 |
| C38D9.2 |
| T03D3.5 |
| *glit-1* |
| *mif-2* |
| *spe-41* |

**Table S3. Gene Ontology terms associated with genes in Cluster 1 among the top 100 genes clustered using historical mutual information.**

| GO term | # in set | # identified | # expected | Enrichment | P value |
|---|---|---|---|---|---|
| regulation of biological process | 4202 | 13 | 4.03 | 3.23 | 3.20E-02 |
| developmental process | 1846 | 10 | 1.77 | 5.65 | 5.10E-03 |

| | | | | | |
|---|---|---|---|---|---|
| cellular component organization | 1830 | 10 | 1.75 | 5.7 | 4.70E-03 |
| cellular component organization or biogenesis | 1982 | 10 | 1.9 | 5.26 | 9.78E-03 |
| anatomical structure development | 1696 | 9 | 1.63 | 5.54 | 2.55E-02 |
| regulation of macromolecule metabolic process | 1646 | 9 | 1.58 | 5.71 | 1.99E-02 |
| regulation of metabolic process | 1745 | 9 | 1.67 | 5.38 | 3.22E-02 |
| cell cycle process | 400 | 9 | 0.38 | 23.48 | 9.93E-08 |
| cell cycle | 517 | 9 | 0.5 | 18.16 | 9.66E-07 |
| negative regulation of biological process | 962 | 8 | 0.92 | 8.68 | 3.55E-03 |
| reproductive process | 803 | 8 | 0.77 | 10.4 | 9.01E-04 |
| sexual reproduction | 425 | 8 | 0.41 | 19.64 | 6.51E-06 |
| cell differentiation | 824 | 8 | 0.79 | 10.13 | 1.10E-03 |
| cellular developmental process | 826 | 8 | 0.79 | 10.11 | 1.12E-03 |
| mitotic cell cycle | 269 | 8 | 0.26 | 31.03 | 1.74E-07 |
| organelle organization | 1091 | 8 | 1.05 | 7.65 | 9.11E-03 |
| negative regulation of cellular process | 854 | 7 | 0.82 | 8.55 | 2.18E-02 |
| mitotic cell cycle process | 233 | 7 | 0.22 | 31.35 | 3.21E-06 |
| embryo development | 516 | 7 | 0.49 | 14.16 | 7.57E-04 |
| regulation of cell cycle | 245 | 6 | 0.23 | 25.55 | 2.01E-04 |
| multicellular organismal reproductive process | 384 | 6 | 0.37 | 16.3 | 2.82E-03 |
| regulation of cell cycle process | 185 | 5 | 0.18 | 28.2 | 1.78E-03 |
| gamete generation | 254 | 5 | 0.24 | 20.54 | 8.45E-03 |
| germ cell development | 176 | 5 | 0.17 | 29.64 | 1.39E-03 |
| cellular process involved in reproduction in multicellular organism | 178 | 5 | 0.17 | 29.31 | 1.47E-03 |
| microtubule cytoskeleton organization | 202 | 5 | 0.19 | 25.83 | 2.74E-03 |
| microtubule-based process | 281 | 5 | 0.27 | 18.57 | 1.38E-02 |
| embryo development ending in birth or egg hatching | 303 | 5 | 0.29 | 17.22 | 2.00E-02 |
| regulatory ncRNA-mediated gene silencing | 109 | 4 | 0.1 | 38.29 | 7.97E-03 |
| regulation of mitotic cell cycle | 94 | 4 | 0.09 | 44.4 | 4.41E-03 |

| | | | | | |
|---|---|---|---|---|---|
| oogenesis | 117 | 4 | 0.11 | 35.67 | 1.06E-02 |
| female gamete generation | 144 | 4 | 0.14 | 28.99 | 2.41E-02 |
| nuclear chromosome segregation | 121 | 4 | 0.12 | 34.49 | 1.21E-02 |
| chromosome segregation | 155 | 4 | 0.15 | 26.93 | 3.22E-02 |
| nuclear division | 166 | 4 | 0.16 | 25.14 | 4.22E-02 |

**Table S4. Potential hypotheses for the function(s) of PIRE proteins without common names.** Function(s) of the known regulators of RNA silencing could be promoted or inhibited by interacting PIRE proteins.

| PIRE | Interactor | Known function(s) of RNA regulator(s) |
|---|---|---|
| PIRE-1/ Y20F4.4 | ADR-2 | A-to-I editing of dsRNA (double-stranded RNA) (8) |
| | HRDE-1 | Argonaute activity (9) |
| PIRE-2/ C08F11.7 | ADR-2 | A-to-I editing of dsRNA (double-stranded RNA) (8) |
| | CSR-1 | Argonaute activity (10) |
| | RDE-3 | poly-UG RNA production (11,12) |
| | SET-25 | histone methyltransferase activity (13,14) |
| PIRE-3/ E01G4.5 | DEPS-1 | germ granule formation and/or RNA silencing (15) |
| | RDE-3 | poly-UG RNA production (11,12) |
| | PGL-1 | mRNA regulation and/or P granule formation (16) |
| | MUT-7 | 3’-5’ exoribonuclease activity (17) |
| | SET-25 | histone methyltransferase activity (13,14) |
| PIRE-4/ F15D4.5 | DEPS-1 | germ granule formation and/or RNA silencing (15) |
| | HRDE-1 | Argonaute activity (9) |
| | PRG-1 | Argonaute activity (18) |
| | RDE-8 | RNA endonuclease and/or mRNA binding activity (19) |
| | RDE-3 | poly-UG RNA production (11,12) |
| PIRE-5/ K02E2.6 | ERI-1 | 3’-5’ exoribonuclease activity (20) |
| | PID-2 | piRNA-mediated silencing and/or Z-granule formation (7) |
| | RDE-8 | RNA endonuclease and/or mRNA binding activity (19) |
| | SET-25 | histone methyltransferase activity (13,14) |
| PIRE-6/ R06C1.4 | RDE-3 | poly-UG RNA production (11,12) |
| | PRG-1 | Argonaute activity (18) |
| PIRE-7/ C38D9.2 | RDE-1 | Argonaute activity (21,22) |
| | PRG-1 | Argonaute activity (18) |
| PIRE-8/ T16G12.4 | CSR-1 | Argonaute activity (10) |
| PIRE-9/ Y47H10A.5 | ADR-2 | A-to-I editing of dsRNA (8) |
| | ERI-1 | 3’-5’ exoribonuclease activity (20) |
| | PID-2 | piRNA-mediated silencing and/or Z-granule formation (7) |
| | MUT-16 | secondary small RNA production and mutator foci formation (23) |
| | SET-25 | histone methyltransferase activity (13,14) |

**Table S5. $r_g$ rank order of frequently identified genes.** Top 25 rank-ordered genes obtained by calculating $r_g$ using 25, 100, or 1000 of the most frequently listed genes among the 398 tables considered in this study. In bold are genes shared with the top 25 identified using the most frequent 1000 genes.

| $r_{25}$ genes | | $r_{100}$ genes | | $r_{1000}$ genes | |
|---|---|---|---|---|---|
| **C55C3.3** | 0.0164 | **C55C3.3** | 0.0164 | **C55C3.3** | 0.0164 |
| ***timm-17B.2*** | 0.0149 | ***timm-17B.2*** | 0.0149 | ***timm-17B.2*** | 0.0149 |
| **Y20F4.4** | 0.0135 | **Y20F4.4** | 0.0135 | **Y20F4.4** | 0.0135 |
| **C08F11.7** | 0.0125 | **C08F11.7** | 0.0125 | **C08F11.7** | 0.0125 |
| **E01G4.5** | 0.0120 | **E01G4.5** | 0.0120 | **E01G4.5** | 0.0120 |
| **ZK402.3** | 0.0119 | **ZK402.3** | 0.0119 | **ZK402.3** | 0.0119 |
| **C09G5.7** | 0.0118 | **C09G5.7** | 0.0118 | **C09G5.7** | 0.0118 |
| ***hrde-1*** | 0.0117 | ***hrde-1*** | 0.0117 | ***hrde-1*** | 0.0117 |
| **C18D4.6** | 0.0117 | **C18D4.6** | 0.0117 | **C18D4.6** | 0.0117 |
| **R06C1.4** | 0.0116 | **R06C1.4** | 0.0116 | **R06C1.4** | 0.0116 |
| **C38D9.2** | 0.0115 | **C38D9.2** | 0.0115 | **C38D9.2** | 0.0115 |
| **F15D4.5** | 0.0115 | **F15D4.5** | 0.0115 | **F15D4.5** | 0.0115 |
| **T16G12.4** | 0.0109 | **Y57G11C.51** | 0.0112 | **Y57G11C.51** | 0.0112 |
| ***fbxb-97*** | 0.0107 | ***pan-1*** | 0.0111 | ***pan-1*** | 0.0111 |
| **W04B5.1** | 0.0103 | ***hil-4*** | 0.0111 | ***hil-4*** | 0.0111 |
| ***spe-41*** | 0.0102 | ***cdk-1*** | 0.0111 | ***cdk-1*** | 0.0111 |
| *scrm-4* | 0.0098 | **T16G12.4** | 0.0109 | **T16G12.4** | 0.0109 |
| F39E9.7 | 0.0098 | ***fbxb-97*** | 0.0107 | ***fbxb-97*** | 0.0107 |
| K02E2.6 | 0.0097 | **F39F10.4** | 0.0106 | **F39F10.4** | 0.0106 |
| W04B5.2 | 0.0096 | **K09H9.7** | 0.0106 | **K09H9.7** | 0.0106 |
| *rnh-1.3* | 0.0095 | ***tbb-2*** | 0.0105 | ***tbb-2*** | 0.0105 |
| *bath-45* | 0.0094 | ***saeg-1*** | 0.0105 | ***saeg-1*** | 0.0105 |
| F58H7.5 | 0.0084 | **W04B5.1** | 0.0103 | **W04B5.1** | 0.0103 |
| SDG-1 | 0.0062 | ***spe-41*** | 0.0102 | ***spe-41*** | 0.0102 |
| W09B7.1 | 0.0032 | ***csr-1*** | 0.0101 | ***csr-1*** | 0.0101 |

**Supplementary Movie Legends**

**Movie S1.** TIMM-17B.2 and ADR-2 with inter-protein predicted aligned error < 5 and distance < 6
**Movie S2.** TIMM-17B.2 and PID-2 with inter-protein predicted aligned error < 5 and distance < 6
**Movie S3.** TIMM-17B.2 and RDE-8 with inter-protein predicted aligned error < 5 and distance < 6
**Movie S4.** TIMM-17B.2 and RDE-3 with inter-protein predicted aligned error < 5 and distance < 6
**Movie S5.** Y20F4.4 and HRDE-1 with inter-protein predicted aligned error < 5 and distance < 6
**Movie S6.** C08F11.7 and ADR-2 with inter-protein predicted aligned error < 5 and distance < 6
**Movie S7.** C08F11.7 and CSR-1 with inter-protein predicted aligned error < 5 and distance < 6
**Movie S8.** C08F11.7 and RDE-3 with inter-protein predicted aligned error < 5 and distance < 6
**Movie S9.** C08F11.7 and SET-25 with inter-protein predicted aligned error < 5 and distance < 6
**Movie S10.** E01G4.5 and PGL-1 with inter-protein predicted aligned error < 5 and distance < 6
**Movie S11.** E01G4.5 and MUT-7 with inter-protein predicted aligned error < 5 and distance < 6

**Movie S12.** E01G4.5 and SET-25 with inter-protein predicted aligned error < 5 and distance < 6

**Movie S13.** F15D4.5 and DEPS-1 with inter-protein predicted aligned error < 5 and distance < 6

**Movie S14.** F15D4.5 and RDE-8 with inter-protein predicted aligned error < 5 and distance < 6

**Movie S15.** F15D4.5 and RDE-3 with inter-protein predicted aligned error < 5 and distance < 6

**Movie S16.** FBXB-97 and RDE-4 with inter-protein predicted aligned error < 5 and distance < 6

**Movie S17.** FBXB-97 and ERI-1 with inter-protein predicted aligned error < 5 and distance < 6

**Movie S18.** FBXB-97 and NRDE-3 with inter-protein predicted aligned error < 5 and distance < 6

**Movie S19.** FBXB-97 and DEPS-1 with inter-protein predicted aligned error < 5 and distance < 6

**Movie S20.** FBXB-97 and PID-2 with inter-protein predicted aligned error < 5 and distance < 6

**Movie S21.** FBXB-97 and RDE-8 with inter-protein predicted aligned error < 5 and distance < 6

**Movie S22.** FBXB-97 and RDE-3 with inter-protein predicted aligned error < 5 and distance < 6

**Movie S23.** K02E2.6 and ERI-1 with inter-protein predicted aligned error < 5 and distance < 6

**Movie S24.** K02E2.6 and PID-2 with inter-protein predicted aligned error < 5 and distance < 6

**Movie S25.** K02E2.6 and RDE-8 with inter-protein predicted aligned error < 5 and distance < 6

**Movie S26.** K02E2.6 and SET-25 with inter-protein predicted aligned error < 5 and distance < 6

**Movie S27.** RNH-1.3 and RDE-1 with inter-protein predicted aligned error < 5 and distance < 6

**Movie S28.** RNH-1.3 and HRDE-2 with inter-protein predicted aligned error < 5 and distance < 6

**Movie S29.** RNH-1.3 and RDE-3 with inter-protein predicted aligned error < 5 and distance < 6

**Movie S30.** RNH-1.3 and SET-25 with inter-protein predicted aligned error < 5 and distance < 6

**Movie S31.** SDG-1 and ADR-2 with inter-protein predicted aligned error < 5 and distance < 6

**Movie S32.** SDG-1 and RDE-8 with inter-protein predicted aligned error < 5 and distance < 6

**Movie S33.** RNH-1.3 and RDE-3 predicted by AlphaFold 2 versus the AlphaFold 3 server

**Movie S34.** FBXB-97 and RDE-3 predicted by AlphaFold 2 versus the AlphaFold 3 server

**Movie S35.** PIRE-4 and RDE-3 predicted by AlphaFold 2 versus the AlphaFold 3 server

**Movie S36.** EGO-1 and W09B7.1 predicted by AlphaFold 2 versus the AlphaFold 3 server

**Movie S37.** Overlay of two models for the RNH-1.3:RDE-3 complex predicted by AlphaFold 2.

**Movie S38.** Overlay of two models for the RNH-1.3:RDE-3 complex predicted by AlphaFold 3.

**Movie S39.** PIRE-1 and RDE-1 with inter-protein predicted aligned error < 5 and distance < 6.

**Movie S40.** DEPS-1 and PIRE-3 with inter-protein predicted aligned error < 5 and distance < 6.

**Movie S41.** RDE-3 and PIRE-3 with inter-protein predicted aligned error < 5 and distance < 6.

**Movie S42.** R06C1.4 and PRG-1 with inter-protein predicted aligned error < 5 and distance < 6.

**Movie S43.** R06C1.4 and RDE-3 with inter-protein predicted aligned error < 5 and distance < 6.

**Movie S44.** C38D9.2 and RDE-1 with inter-protein predicted aligned error < 5 and distance < 6.

**Movie S45.** C38D9.2 and PRG-1 with inter-protein predicted aligned error < 5 and distance < 6.

**Movie S46.** PRG-1 and PIRE-4 with inter-protein predicted aligned error < 5 and distance < 6.

**Movie S47.** HRDE-1 and PIRE-4 with inter-protein predicted aligned error < 5 and distance < 6.

**Movie S48.** T16G12.4 and CSR-1 with inter-protein predicted aligned error < 5 and distance < 6.

**Movie S49.** ADR-2 and the longest peptide that could be encoded by F39E9.7 with inter-protein predicted aligned error < 5 and distance < 6.

**Movie S50.** CSR-1 and the longest peptide that could be encoded by F39E9.7 with inter-protein predicted aligned error < 5 and distance < 6.

**Movie S51.** RDE-8 and the longest peptide that could be encoded by F39E9.7 with inter-protein predicted aligned error < 5 and distance < 6.

**Movie S52.** PRG-1 and the longest peptide that could be encoded by W04B5.2 with inter-protein predicted aligned error < 5 and distance < 6.

**Movie S53.** RDE-8 and the longest peptide that could be encoded by W04B5.2 with inter-protein predicted aligned error < 5 and distance < 6.

**Movie S54.** ADR-2 and the longest peptide that could be encoded by ZK402.3 with inter-protein predicted aligned error < 5 and distance < 6.
**Movie S55.** ADR-2 and STAU-1 with inter-protein predicted aligned error < 5 and distance < 6.
**Movie S56.** RDE-1 and STAU-1 with inter-protein predicted aligned error < 5 and distance < 6.
**Movie S57.** PRG-1 and STAU-1 with inter-protein predicted aligned error < 5 and distance < 6.
**Movie S58.** ALG-2 and STAU-1 with inter-protein predicted aligned error < 5 and distance < 6.
**Movie S59.** CSR-1 and STAU-1 with inter-protein predicted aligned error < 5 and distance < 6.
**Movie S60.** RDE-8 and STAU-1 with inter-protein predicted aligned error < 5 and distance < 6.
**Movie S61.** ADR-2 and RDE-3 with inter-protein predicted aligned error < 5 and distance < 6.
**Movie S62.** CSR-1 and RDE-3 with inter-protein predicted aligned error < 5 and distance < 6.
**Movie S63.** DEPS-1 and RDE-3 with inter-protein predicted aligned error < 5 and distance < 6.
**Movie S64.** ERGO-1 and RDE-3 with inter-protein predicted aligned error < 5 and distance < 6.
**Movie S65.** MUT-16 and RDE-3 with inter-protein predicted aligned error < 5 and distance < 6.
**Movie S66.** NRDE-3 and RDE-3 with inter-protein predicted aligned error < 5 and distance < 6.
**Movie S67.** PID-2 and RDE-3 with inter-protein predicted aligned error < 5 and distance < 6.
**Movie S68.** PIK-1 and RDE-3 with inter-protein predicted aligned error < 5 and distance < 6.
**Movie S69.** PIR-1 and RDE-3 with inter-protein predicted aligned error < 5 and distance < 6.
**Movie S70.** RDE-4 and RDE-3 with inter-protein predicted aligned error < 5 and distance < 6.
**Movie S71.** RDE-8 and RDE-3 with inter-protein predicted aligned error < 5 and distance < 6.
**Movie S72.** RDE-10 and RDE-3 with inter-protein predicted aligned error < 5 and distance < 6.
**Movie S73.** PID-5 and PID-2 with inter-protein predicted aligned error < 5 and distance < 6.
**Movie S74.** PID-4 and PID-2 with inter-protein predicted aligned error < 5 and distance < 6.
**Movie S75.** KIN-19 and PID-2 with inter-protein predicted aligned error < 5 and distance < 6.
**Movie S76.** PAR-5 and PID-2 with inter-protein predicted aligned error < 5 and distance < 6.
**Movie S77.** T07C4.3 and PID-2 with inter-protein predicted aligned error < 5 and distance < 6.
**Movie S78.** Overlay of models for the PAR-5:PID-2 complex predicted by AlphaFold 2 and AlphaFold 3.
**Movie S79.** Overlay of models for the GOA-1:DGK-1 complexes predicted by AlphaFold 2 and AlphaFold 3.
**Movie S80.** CSR-1 and MCM-7 with inter-protein predicted aligned error < 5 and distance < 6.
**Movie S81.** ADR-2 and PAN-1 with inter-protein predicted aligned error < 5 and distance < 6.
**Movie S82.** ERI-1 and PAN-1 with inter-protein predicted aligned error < 5 and distance < 6.
**Movie S83.** HRDE-1 and PAN-1 with inter-protein predicted aligned error < 5 and distance < 6.
**Movie S84.** HRDE-2 and PAN-1 with inter-protein predicted aligned error < 5 and distance < 6.
**Movie S85.** MUT-7 and PAN-1 with inter-protein predicted aligned error < 5 and distance < 6.
**Movie S86.** RDE-1 and CEY-2 with inter-protein predicted aligned error < 5 and distance < 6.
**Movie S87.** RDE-8 and CEY-2 with inter-protein predicted aligned error < 5 and distance < 6.
**Movie S88.** RDE-3 and CEY-2 with inter-protein predicted aligned error < 5 and distance < 6.
**Movie S89.** Y47H10A.5 and ADR-2 with inter-protein predicted aligned error < 5 and distance < 6
**Movie S90.** Y47H10A.5 and ERI-1 with inter-protein predicted aligned error < 5 and distance < 6.
**Movie S91.** Y47H10A.5 and MUT-16 with inter-protein predicted aligned error < 5 and distance < 6
**Movie S92.** Y47H10A.5 and PID-2 with inter-protein predicted aligned error < 5 and distance < 6
**Movie S93.** Y47H10A.5 and SET-25 with inter-protein predicted aligned error < 5 and distance < 6
**Movie S94.** Simulation illustrating the growth of networks through preferential attachment (Screen capture of ‘Preferential Attachment Simple’ from NetLogo model library).
